# Consistent accumulation of transposable elements in species of the Hawaiian *Tetragnatha* spiny-leg adaptive radiation across the archipelago chronosequence

**DOI:** 10.1101/2024.01.03.574070

**Authors:** Heidi Yang, Clément Goubert, Darko D. Cotoras, Natalie R Graham, José Cerca, Rosemary G. Gillespie

## Abstract

The ecological and phenotypic diversity observed in oceanic island radiations presents an evolutionary paradox: a high level of genetic variation is typically required for diversification, but species colonizing a new island typically suffer from founder effects. This reduction in population size leads to a reduction in genetic diversity, which ultimately results in a reduction in the efficiency of natural selection. Then, what is the source of genetic variation which acts as the raw material for ecological and phenotypic diversification in oceanic archipelagos? Transposable elements (TEs) are mobile genetic elements that have been linked to the generation of genetic diversity, and evidence suggests that TE activity and accumulation along the genome can result from reductions in population size. Here, we use the Hawaiian spiny-leg spider radiation (*Tetragnatha*) to test whether TE accumulation increases due to demographic processes associated with island colonization. We sequenced and quantified TEs in 23 individuals from the spiny-leg radiation and 4 individuals from its sister radiation, the Hawaiian web-building *Tetragnatha*. Our results show that founder effects resulting from colonization of new islands have not resulted in TE accumulation over evolutionary time. Specifically, we found no evidence for increase in abundance of specific TE superfamilies, nor an accumulation of ‘young TEs’ in lineages which have recently colonized a new island or are present in islands with active volcanoes. We also found that the DNA/hAT transposon superfamily is by far the most abundant TE superfamily in the *Tetragnatha* radiation. This work shows that TE abundance has remained constant for the spiny-leg radiation across the archipelago chronosequence, and TE accumulation is not affected by population oscillations associated with island colonization events. Therefore, despite their known role in the generation of genetic diversity, TE activity does not appear to be the mechanism to explain the evolutionary paradox of the insular *Tetragnatha* spiny-leg radiation.

## Introduction

Adaptive radiation, the rapid diversification of a lineage into a wide range of ecological niches, provides a clear link between ecology and evolution (Schluter, 2000; Gillespie *et al*., 2020). Many adaptive radiations are characterized by remarkable phenotypic and ecological variation, but the mechanisms initiating the process of diversification have proven to be difficult to study. A notable paradox is the high number of adaptive radiations observed on oceanic archipelagos, despite the fact that island colonization itself is associated with a drastic reduction of genetic diversity. The species’ arrival to a remote oceanic island entails a strong founder event, which inevitably depletes genetic variation (Cerca *et al*., 2023b). Additionally, insular lineages are often subject to environmental disturbances and stochastic events that lead to local extinctions and population reductions (Frankham, 1997). A reduction in effective population size (N_e_) will lead to a reduction of its genetic diversity, and ultimately in a reduction of the efficiency of natural selection (Charlesworth, 2009). Thus, identifying the source of genetic variation which acts as the raw material for ecological and phenotypic diversification in oceanic archipelagos remains an active question for evolutionary biologists.

Transposable elements (TEs) are mobile genetic sequences that can generate a variety of mutations, including changes in gene coding regions, cis-regulatory elements, 3D chromatin structure, affecting gene regulation, and influencing overall genome sizes (Chénais *et al*., 2012; Belyayev, 2014; Chuong, Elde, & Feschotte, 2017; Fambrini *et al*., 2020; Choudhary *et al*., 2023). For example, a TE insertion has been implicated in the origin and evolution of the industrial melanism ecotype in the peppered moth, a textbook example of rapid adaptation (Hof *et al*., 2016). Accordingly, TE activity has been associated with the rise of novel traits, and diversification dynamics in the context of adaptive radiations. In African cichlids, a TE insertion in the *cis*-regulatory region of a pigmentation gene led to the evolution of egg-spots (Santos *et al*., 2014). Accumulations of TEs in *Hox* genes have been documented in the *Anolis* lizard radiation (Feiner, 2016), and in *Heliconius* butterflies have higher TE abundance when compared to their outgroups (Feiner, 2016; Cicconardi *et al*., 2023). A high accumulation of TEs has been found in African cichlid fishes relative to outgroup taxa (Brawand *et al*., 2014), although recent evidence contradicts this (Ronco et al. 2021). Given the expected lack of genetic variation in populations present on recently colonized areas and that TEs are known facilitators of rapid adaptive evolution, TEs could be hypothesized as candidates in facilitating the generation of genetic variation required for the adaptive radiation to initiate and unfold in insular environments (Oliver & Greene, 2012; Casacuberta & González, 2013; Brawand *et al*., 2014; Ricci *et al*., 2018; Schrader & Schmitz, 2019; Ronco *et al*., 2021).

While TE insertions are thought to be mostly deleterious and thus removed from the population (Kidwell & Lisch, 1997), the genomic accumulation and subsequent contribution to genetic diversity of TEs has been linked to demographic oscillations. For example, reductions in N_e_ alter the efficiency of selection, thereby decreasing the efficiency of purifying selection to remove deleterious TE insertions (Blass, Bell, & Boissinot, 2012; Tollis & Boissinot, 2013; Xue *et al*., 2018; Bourgeois & Boissinot, 2019). Indeed, differential fixation of TEs in *Drosophila subobscura* (García Guerreiro *et al*., 2008) and *Arabidopsis lyrata* (Lockton, Ross-Ibarra, & Gaut, 2008) suggest that oscillations in demography, such as bottlenecks, can increase TE accumulation patterns along genomes due to weak selection (Bourgeois & Boissinot, 2019). However, there is nuance in demography’s role; for example, purifying selection did not constrain the spread of L1 retrotransposons in *Anolis* lizards (Tollis & Boissinot, 2013). Overall, demographic factors likely play a crucial role in determining the likelihood of TEs reaching fixation, with increased accumulation of TEs leading to novel mutations, which could underlie phenotypic and ecological diversity. In this regard, island oceanic radiations are perfect candidates to study the link between demography and TE accumulation patterns. The known age and sequential arrangements of the islands provide an explicit temporal framework and the strong changes in N_e_ as a result of new island colonization could act as a means of TE accumulation.

The Hawaiian island archipelago offers a unique opportunity to examine the dynamics of accumulation of TEs over the course of adaptive radiation, since the archipelago comprises a geological chronosequence of volcanic islands –– resembling ‘evolution on a conveyor belt’ (Funk and Wagner 1995, Fleischer et al. 1998). Most lineages in the archipelago have colonized older islands, and progressed down the island chain as newer islands have formed (Shaw & Gillespie, 2016). The *Tetragnatha* spiny-leg spider radiation follows this colonization pattern from older to younger islands, which consists of 17 species that divergently evolved into one of four ecomorph types: “maroon”, “green”, “large brown”, and “small brown” (Gillespie, 1991, 2004; Roderick & Gillespie, 1998; Cerca *et al*., 2023a). Since the radiation unfolds from older to younger islands, we can test whether the more recent colonization events on the younger islands correspond to an increase in TE accumulation. During sequential island colonization, a reduction of population size, and associated reduction in N_e_, occurs due to dispersal limitation across open water to newly forming volcanoes. Once early colonists arrive, they encounter highly heterogeneous landscapes, where lava flows frequently fragment Hawaiian rainforests and result in the formation of small, isolated, and transient pockets of forest (kīpuka) that further alter patterns of demography and speciation. Indeed, the habitat fragmentation has been proposed as a mechanism in driving diversification by creating a metapopulation dynamic with intermittent events of isolation and admixture (Carson & Templeton, 1984; Vandergast, Gillespie, & Roderick, 2004; Roderick *et al*., 2012).

Using volcano age as a proxy for substrate age and colonization history, we test the hypothesis that population reductions associated with new island colonizations––and intra-island volcano colonizations––lead to an increase of TE accumulation in spiny leg *Tetragnatha*. We then expect that species on the youngest volcanoes will have the highest overall abundance of TEs and highest proportion of young TEs, reflecting an increase in TE accumulation. To this end, we whole-genome resequenced 23 spiny-leg *Tetragnatha* individuals from the Hawaiian archipelago and characterized the TE accumulation within the genomes, by estimating abundance of specific TE superfamilies, exploring the differential accumulation of young TEs, and correlating volcano age with TE accumulation.

## Methods

### Data set

The collection and sequencing of the dataset has been described by Cerca et al. (2023*b*) and is presented in Supplementary Table 1. Briefly, we selected a total of 27 individuals, representing 16 species in the spiny-leg *Tetragnatha* radiation, and included 11 additional specimens of *T. anuenue*, *T. brevignatha*, *T. kamakou*, and *T. quasimodo* from different volcanic communities to account for volcano-specific differences in TE accumulation (Figure 1a). The oldest volcano on which *Tetragnatha* is present is the dormant shield volcano on Kauaʻi (ca. 5 Mya), and the youngest are the currently active volcanoes on Hawaiʻi: Mauna Loa (emergent lava flow to 700,000-1 Mya) and Kīlauea (emergent lava flow to 210,000-280,000 years old) (Figure 1b; Table 1). We complemented this dataset with four species of a sister lineage, the web-builder clade, including *T. maka* (Kauaʻi), *T. acuta* (East Maui), *T. filiciphilia* (East Maui), *T. stelarobusta* (East Maui) (Table 1).

**Figure 1a.**
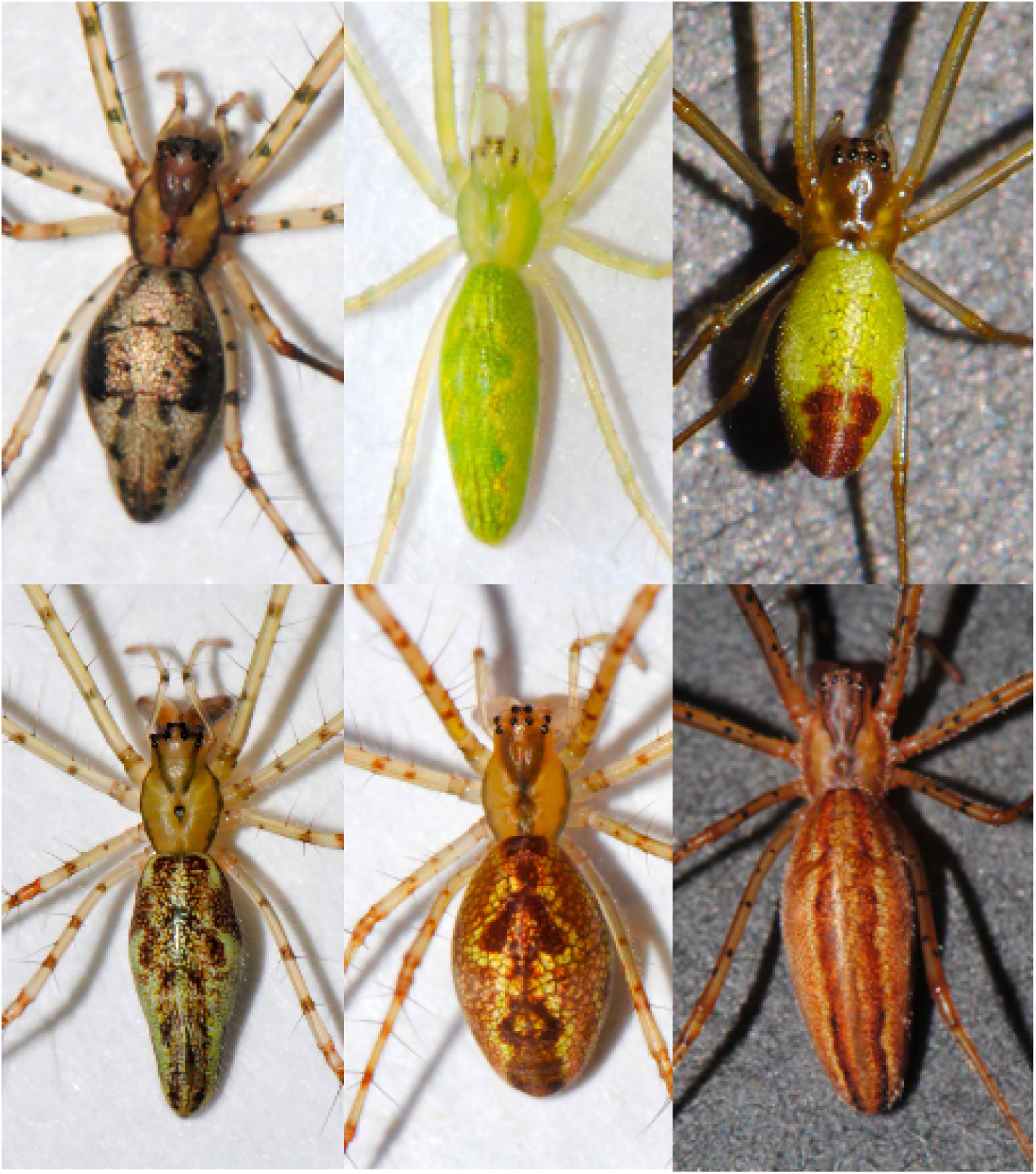
Representative photographs of *Tetragnatha* spiny-leg species, including *Tetragnatha mohihi* (top left, small brown, spiny-leg clade), T. tantalus (top center, green, spiny-leg clade), *T. filiciphilia* (top right, web-building clade), *T. pilosa* (bottom left, big brown, spiny-leg clade), *T. perreirai* (bottom center, maroon, spiny-leg clade), *T. stelarobusta* (bottom right, web-building clade).

**Figure 1b.**
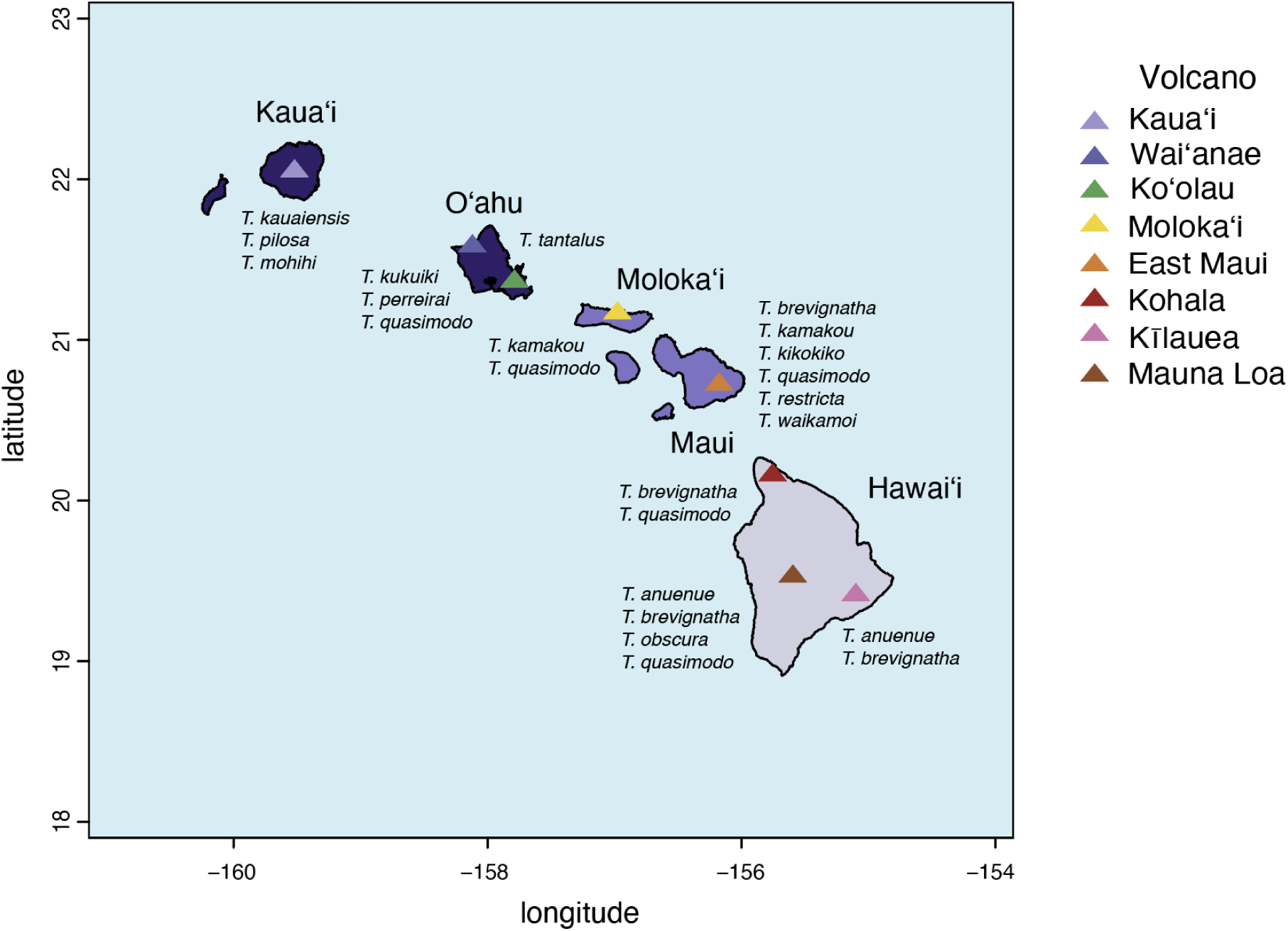
Map of *Tetragnatha* spiny-leg individuals. Islands are colored by old (dark purple), middle-aged (medium purple), and young (light purple) with regard to their relative substrate age. Each island is in some cases the composite of multiple volcanos, each indicated by colored triangles in the key. Several *Tetragnatha* species included in the study are single island endemics. *Tetragnatha anuenue* (Kīlauea, Mauna Loa), *T. brevignatha* (Kohala, Kīlauea), & *T. quasimodo* (Waiʻanae, Molokaʻi, East Maui, Mauna Loa) are included in multiple volcanoes.

**Table 1.**
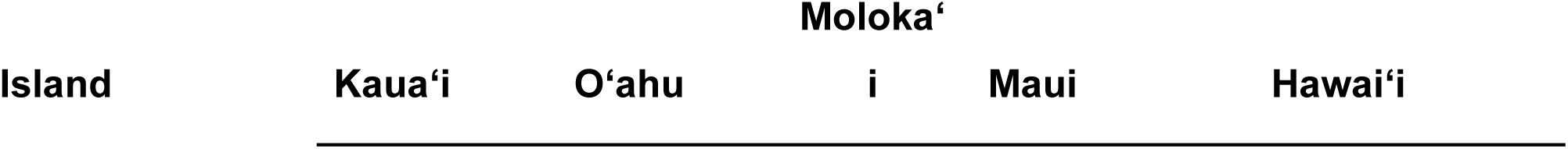
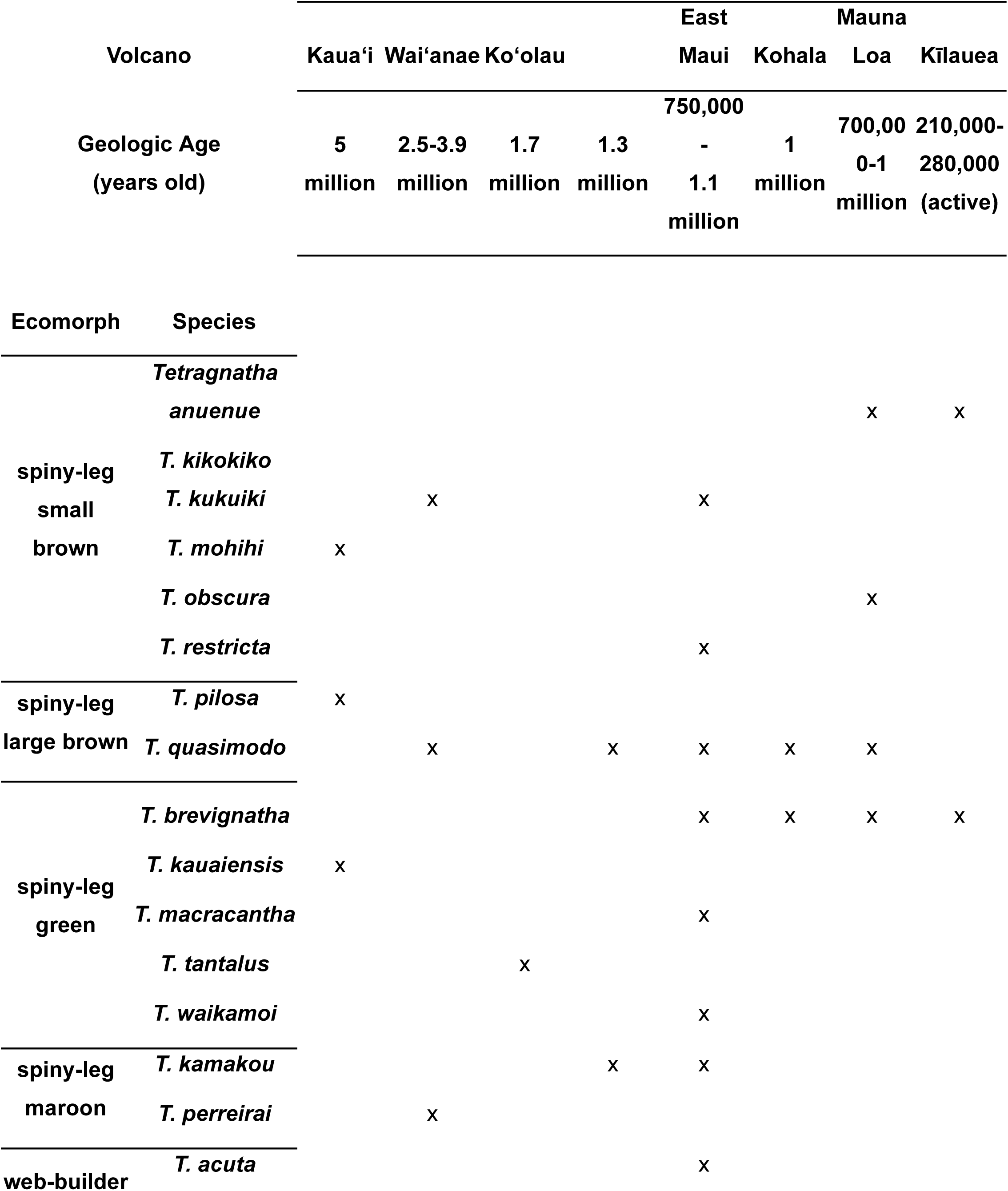
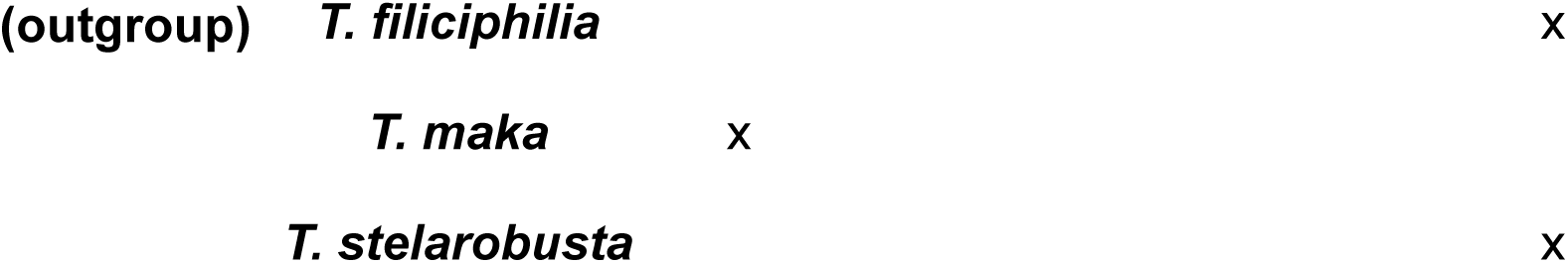
Data set of *Tetragnatha* specimens. The following table shows which volcano and ecomorph type is associated with each specimen. There are 23 total spiny-leg specimens and 4 web-builders.

### Phylogeny of the spiny-leg clade

To compare TEs across individuals in our sample, we started by constructing a molecular phylogeny of the species in our data set. We examined this since there is evidence of hybridization in the spiny-leg radiation (Cerca *et al*., 2023a), which can lead to topological differences in different datasets. First we cleaned the raw Illumina data by identifying and removing adapters using AdapterRemoval *v2.3.2* (Schubert et al. 2016). We used *skmer v3.2.1.* (Marçais and Kingsford 2011; Ondov et al. 2016; Sarmashghi et al. 2019; Rachtman et al. 2022), a *k-mer* based method that estimates genetic distance between genome skims, or low-coverage sets of reads. Then we estimated the pairwise evolutionary distances between the 27 individuals (Sarmashghi et al. 2019). This involved running ‘*skmer reference*’, followed by ‘*skmer subsample*’ to create 100 subsamples of the library as detailed in best practices. Finally, we used ‘*skmer correct*’ to obtain corrected distance matrices of the subsample replicates. We then used *fastme v2.1.5* (Lefort et al. 2015) to obtain a neighbor-joining tree, and we rooted the tree with the sister taxa web-building clade (*T. acuta*, *T. filiciphilia*, *T. maka*, *T. stelarobusta*).

### Classification and Quantification of TEs

To classify and quantify TEs, we used dnaPipeTE, a pipeline designed for annotating, classifying, and quantifying TEs in low-coverage genome samples (< 1x coverage) (Goubert *et al*., 2015). DnaPipeTE also identifies repeat elements such as low-copy repeats and satellite repeats, which we included in our analyses. Before running dnaPipeTE, we generated a *de novo* library of repeats for the *Tetragnatha kauaiensis* genome assembly produced by Cerca et al. (2021) using Repeat Modeler v2.0.2 (Flynn *et al*., 2020)). In addition, we obtained arthropod-specific repeats consensus from Repbase (version 2014-01-31) (Jurka *et al*., 2005). We then concatenated the *de novo* library and the RepBase-based database to obtain a final TE/repeat library.

We ran dnaPipeTE for each individual using the final TE/repeat library and specifying-genome_size 1100000000 –genome_coverage 0.25 –sample_number 2 for all individuals. DnaPipeTE produced TE and repeat contigs (complete or partial assembly of the average (consensus) sequence of the recovered TE families and repeats), classified according to the annotation provided in the TE/repeat library. Using the divergence between short-reads and these consensus contigs, dnaPipeTE estimated repeat landscapes, which approximate the relative age of TE families and allowed us to classify TEs as young/old based on % divergence. We used TE classification at the order, superfamily, and family level from dnaPipeTE outputs.

### Comparative Statistical Analyses

We used Pearson’s correlation test to determine the relationship between volcano age and genome proportion as calculated by dnaPipeTE across the lineage. We ran additional statistical tests for each major TE or repeat element group (DNA, LINE, low-complexity repeat, LTR, simple repeat, satellite repeat, and SINE) to determine if there are element-specific trends in TE/repeat accumulation. Specifically, we grouped individuals by volcano and averaged their TE abundances to perform comparative tests. We used the Kruskal-Wallis test as a nonparametric way to test for significant differences among groups for both genomic TE proportions of each element and percentage of young TEs. We considered young TEs to be those less than 5% divergent as calculated by dnaPipeTE, and calculated proportions of young TEs out of total repetitive element content. If TEs are more active in more recently formed islands, we expected to see higher TE accumulation on individuals from younger volcanoes. As we found the DNA/hAT transposons to be the most abundant superfamily, we additionally tested if there are any significant differences between volcanic communities using a Kruskal-Wallis test for this subset of the data independently, using the same volcano level groupings and dependant variables as the whole TE analyses.

## Results

### TE abundance and diversity in Tetragnatha spiny-legs

Repetitive elements, which include TEs, satellite repeats, and low-complexity repeats, represent 16.15% of the genome on average across both the Hawaiian *Tetragnatha* spiny leg clade (Figure 2). From these, TEs alone make up on average 15.38% of the genome in the spiny legs. There is little variation across genomes in terms of the absolute TE content and the relative percentage of TEs in different species (Figure 2). After filtering for TEs with classifications at the superfamily level, DNA transposons make up the highest percentage, specifically DNA/hAT transposons which on average comprised 30.41% of classified TEs. The next most abundant TE superfamilies are LTR/Gypsy (6.71%), DNA/Academ (6.10%), RC/Helitron (4.95%), DNA/CMC (4.63%), and LINE/1 (4.63%) (Figure 2). We find little variation in the accumulation of TE superfamilies across individuals in the spiny-leg radiation and in the sister orb-weaver radiation (Figure 3). Although DNA/hAT transposons are the most abundant of all TEs, we did not find significant differences in abundance between volcanic communities (Kruskal-Wallis chi-squared, *p*-value = 0.4335).

**Figure 2.**
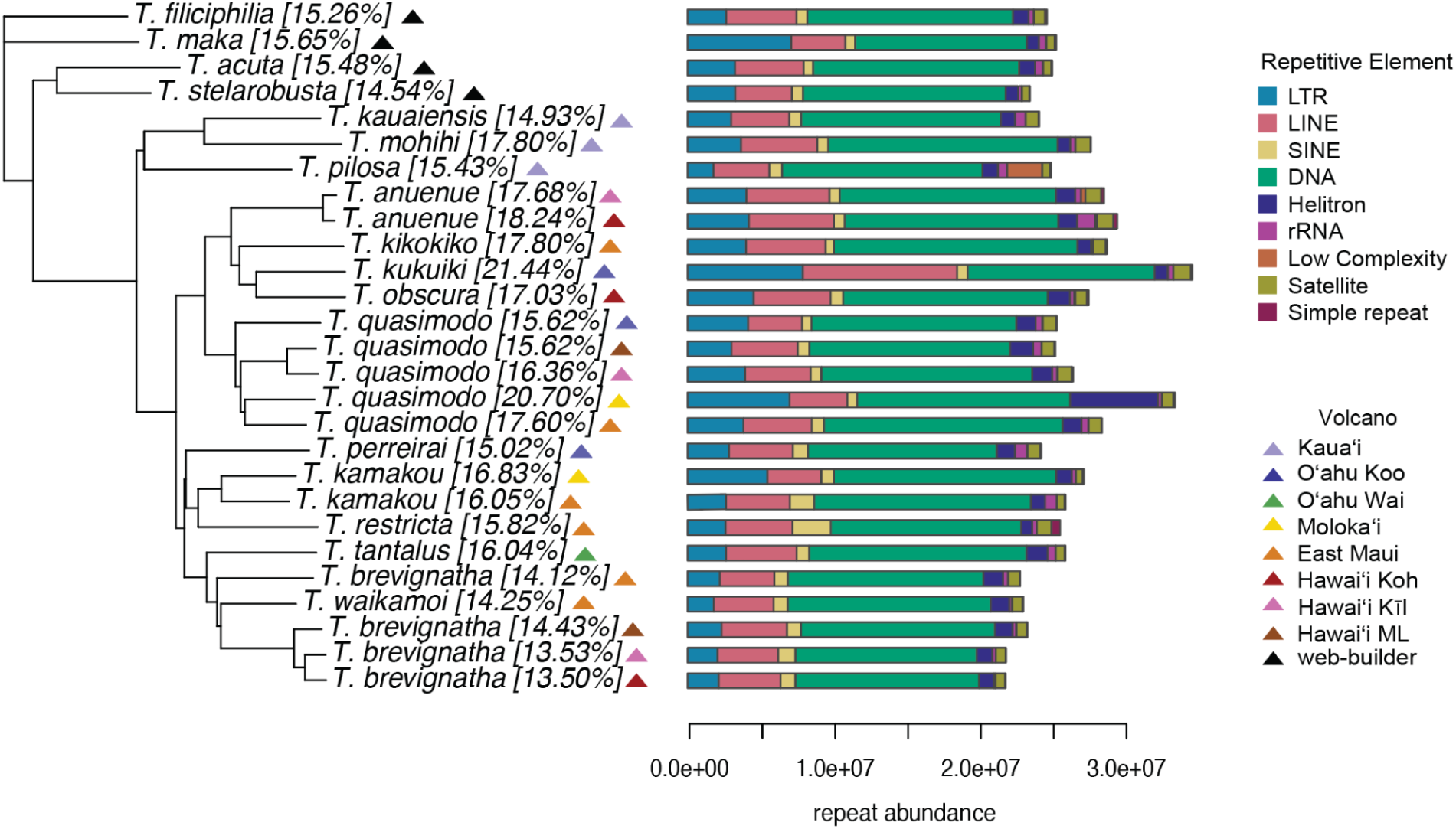
Abundance of repetitive elements across Hawaiian *Tetragnatha* spiny-leg phylogeny. DNA transposons comprise the majority of repetitive elements in all individuals. Percentage of genomic repeat content are in brackets next to each individual. *T. kukuiki* has the highest proportion of genomic repeat content (21.44%) and *T. brevignatha* 067 has the lowest (13.50%). The units of repeat abundance are in basepairs.

**Figure 3.**
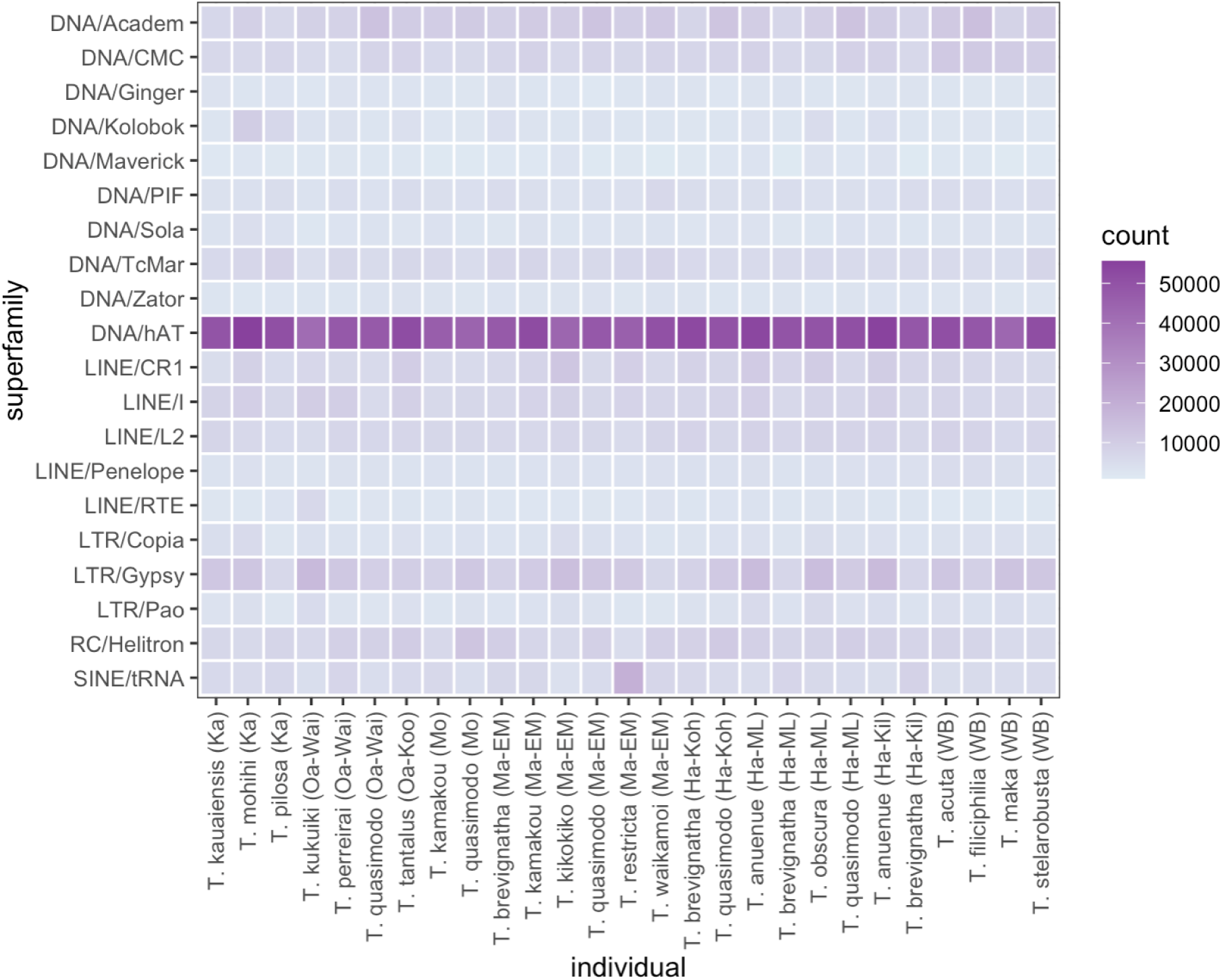
**Accumulation of TE superfamilies in Hawaiian *Tetragnatha***. The X axis shows different species, and the Y axis shows the accumulation of different TE superfamilies. The 20 most abundant superfamilies are included.

### TE Age Distributions

The repeat landscape shows that most TEs have low divergence when aligned to the consensus (Figure 4). All individuals have two peaks of varying sizes: the largest peak is close to 0% divergence and a smaller peak is around 5% divergence (all individual repeat landscape plots are given in Supplementary Figure 1). *T. restricta* from East Maui (Supplementary Figure 1) has a unique distribution pattern compared to the other spiny leg species, with a much higher representation of LINEs, LTRs, and SINEs in the first age peak.

**Figure 4.**
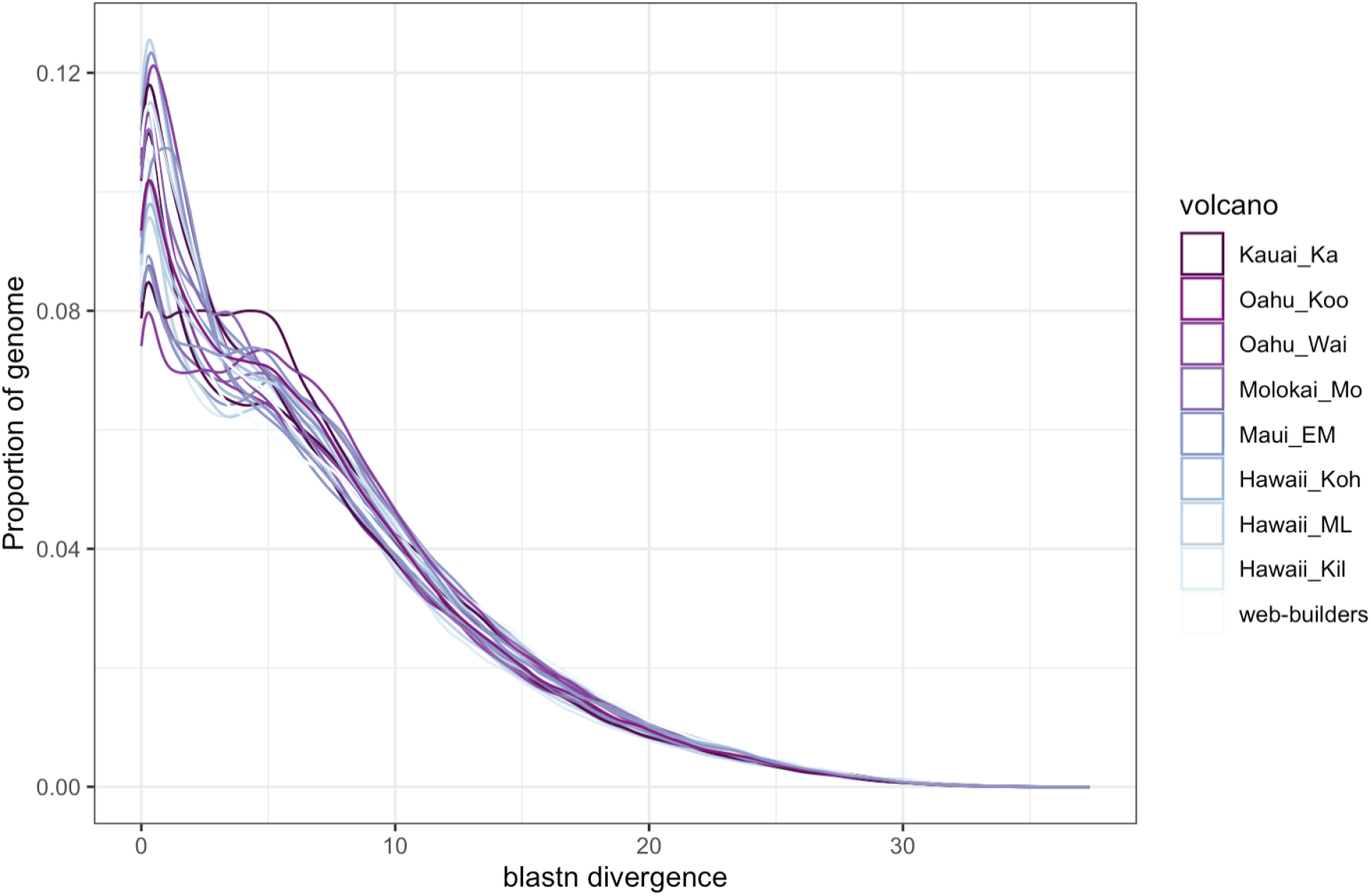
Repeat accumulation plot in *Tetragnatha* spiny-legs. Grouped by volcano, *Tetragnatha* individuals across the spiny-leg lineage and outgroup web-builder lineage exhibit similar patterns of TE age. All individuals exhibit similar patterns of a peak of TEs at <5% blastn divergence, a smaller peak of TEs at 5% divergence, and a steady decline of TEs at higher divergence levels.

The percentage of young TEs (i.e. TEs with an average of <5% divergence between reads and dnaPipeTE consensus) ranged between 40.85% and 50.82%, with an average of 45.63%. We found no significant difference in young TE percentages across volcanoes (Kruskal-Wallis chi-squared, *p*-value > 0.05) (Figure 4).

### Statistical Analyses

The correlation between volcanic age and total repeat abundance was not significant (r = 0.002, *p*-value = 0.9928), indicating no linear relationship between the two (Figure S5). Abundance of specific repeat elements did not differ significantly between volcanoes (Table 2a, Figure S4), with the exception of satellite repeat abundance which differed significantly between the spiny leg and the web-building clade (*p*-value = 0.037) (Table 2b). Finally, the abundance of young TEs did not differ significantly between species from different volcanoes (Kruskal-Wallis chi-squared= 1.164, *p*-value = 0.992).

**Table 2.** Outcome of Kruskal-Wallis tests based on volcano for spiny leg individuals and (b) Mann-Whitney tests for comparing spiny leg and web-builder individuals for each repetitive element type.

| Element | Kruskal-Wallis chi-squared | *p*-value |
| --- | --- | --- |
| DNA | 7.149 | 0.414 |
| Helitron | 7.842 | 0.346 |
| Low-complexity | 2.853 | 0.898 |
| LINE | 6.022 | 0.537 |
| LTR | 9.188 | 0.239 |
| rRNA | 5.465 | 0.603 |
| Satellite | 6.219 | 0.514 |
| SINE | 2.602 | 0.919 |
| Simple repeat | 4.986 | 0.662 |

## Discussion

This study investigated the relationship between transposable element (TE) accumulation on the genome and species/population age in the *Tetragnatha* spiny-leg radiation, benefiting from the Hawaiian chronosequence. We hypothesized that the demographic oscillations associated with early island colonization events would trigger an increase of TE accumulation along the genome. We postulated that this mechanism will explain an increase in genetic variability despite its previous reduction due to founder effects. To test this hypothesis, we analyzed the repeat abundance over a phylogenetic backbone (Figure 2), accumulation of repeats per superfamily (Figure 3), and unveiled the age of TEs (Figures 4-5), with a specific focus on young TEs (Figure 5). We formally tested the accumulation of repeat content (overall and different repeat element group) with volcanic age, finding no significant relationships nor any correlation. If strong population oscillations would have led to an increase in TE activity, we expected species from younger volcanoes to have the highest accumulation of TEs overall as well as young TEs. However, we observed no such trends and instead found little variation across the radiation, suggesting a consistent accumulation of TEs in all individuals, regardless of volcanic community age.

**Figure 5.**
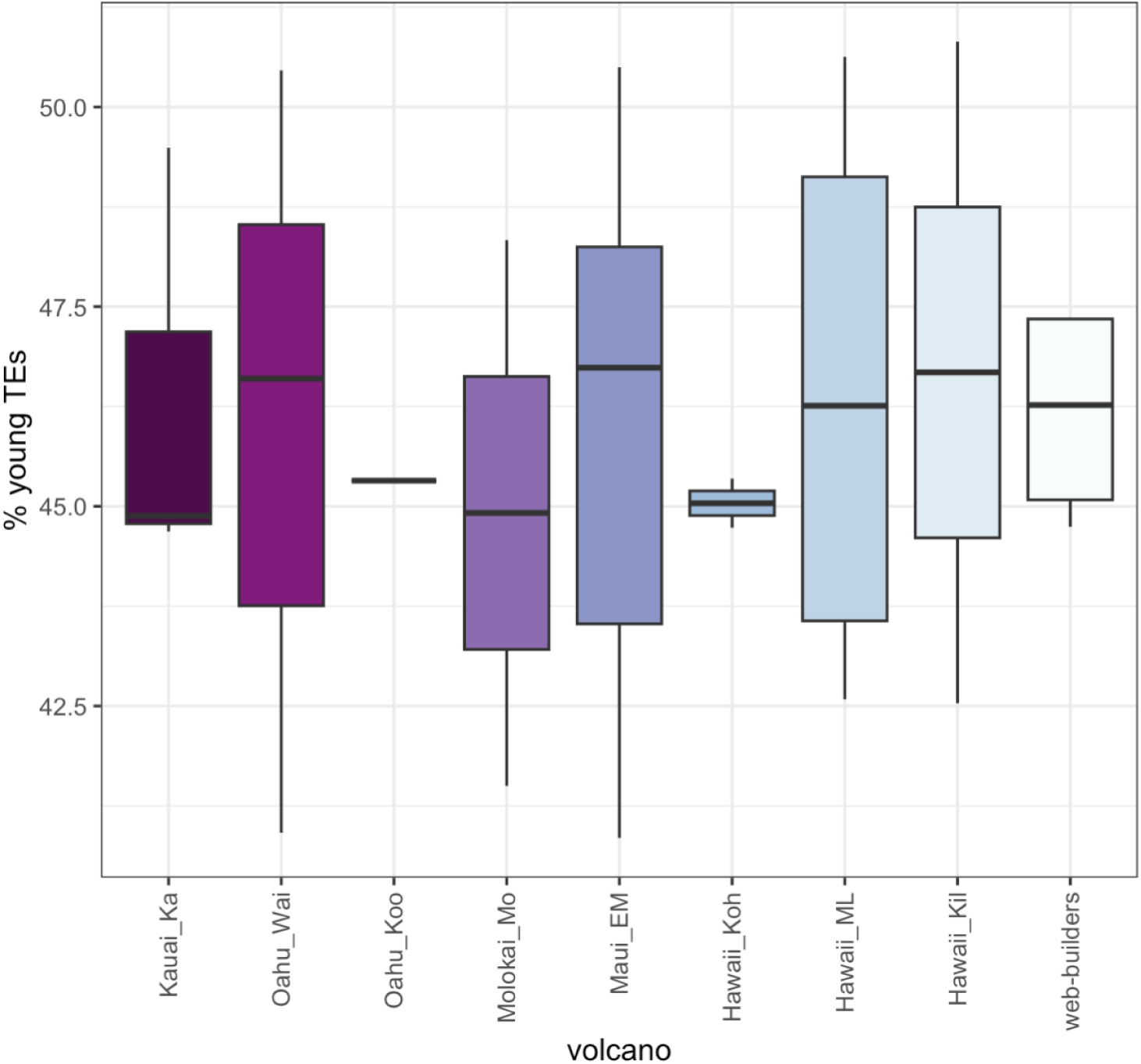
Percentage of young TEs across volcanoes. Percentage of TEs with less than 5% divergence from their consensus sequence.

### No accumulation of overall TEs after early island colonization events

In the *Tetragnatha* spiny-leg adaptive radiation, the accumulation of TEs was not affected by island colonization. The trajectory of island colonization can be analyzed in a comparative phylogenetic setting (Figure 2; Table 1), and this phylogeny is topologically consistent with previous *Tetragnatha* phylogenies (Cerca *et al*., 2023a).

Regardless of island age, genomic repeat content remained fairly consistent across all spiny-leg individuals. For instance, lineages from the oldest island, Kauaʻi, have an average of 15.82% of repeats, which is close to the average of 16.15% of the entire spiny-leg dataset. Species on the volcanically active island of Hawaiʻi have an average of 15.8% genomic repeat content. Similarly, species that were sampled on multiple islands had no particular differences. Additionally, individuals from the same species occurring in different volcanoes had some variation in repeat content, but this is not correlated to the age of the island. The two individuals of *T. anuenue* from Hawaiʻi have 17.68 – 18.24% of their genome composed by repeats, the three *T. brevignatha* also from the island of Hawaiʻi ranges between 13.5 – 14.43%, and the two *T. kamakou* from East Maui and Molokaʻi ranged between 16.05 – 16.83%. There was larger variation in *T. quasimodo* individuals, which we sampled from 4 different islands, which ranges from 15.62 – 20.7%. These results suggest that the founder effects resulting from the colonization of a novel island do not impact the overall repeat content.

The lack of differences in overall accumulation of TEs as a result of population oscillations following a founder event suggests that demography does not greatly influence TE content on *Tetragnatha* genomes. The evidence for the role of demography in promoting TE diversity along genomes has been mixed (reviewed in Bourgeois & Boissinot 2019), and it is possible that oscillations in demography related to early island colonization increases the number of active TE transcripts or the transcription rates of TEs, instead of increasing TE accumulation (Picot *et al*., 2008; García Guerreiro *et al*., 2008; Blass *et al*., 2012; Tollis & Boissinot, 2013). In any case, we expected that the founder events, together with the evidence for small population sizes in insular species would result in TE accumulation in *Tetragnatha*, as natural selection operates less efficiently in small population and would ultimately increase overall TE content. Given our results, we reject the hypothesis that a reduction of population sizes are compromising the action of purifying selection in removing TEs along genomes (Blass et al. 2012; Tollis and Boissinot 2013; Xue et al. 2018; Bourgeois and Boissinot 2019). We recommend that future works should focus on quantifying both transcripts and genomic evolution of TEs at the family level, to obtain a full picture of repeat evolution.

Volcanic activity across islands has been shown to be an important mechanism in driving population structuring of Hawaiian lineages (Wagner & Funk, 1995; Roderick *et al*., 2012), which can create opportunities for geographic isolation and secondary contact, ultimately catalyzing speciation and adaptation (Schluter, 2000; Cotoras *et al*., 2018; Marques, Meier, & Seehausen, 2019; Cerca *et al*., 2023b). Considering the oscillations resulting from the reduction of populations into isolated paths of forests (kīpuka), together with the stress imposed on populations and individuals following volcanic activities (Craddock, 2016), we hypothesized that lineages inhabiting younger islands and volcanoes would have a higher accumulation of TEs. However, we reject these hypotheses, as we did not find significant correlations of age-TE accumulation, nor differences in genomic proportions of specific repeat elements (Table 2) and young TEs (Figures 4, 5).

### No accumulation of specific TEs after early island colonization events

The analyses of TE order (e.g. LTRs, DNA; Figure 2) and specific superfamilies (Figure 3) show little TE diversity across individuals. In theory, one of the most common observations in TE accumulation is the release and accumulation of only a single TE family (Hawkins *et al*., 2006). For instance, genome deregulation as a result of stress, hybridization, or other evolutionary events could cause one TE repression mechanism (i.e. targeted TE methylation, smallRNA, piRNA in TEs, histone marks modification) to be less efficient, leading to the release and expansion of a specific family (Slotkin & Martienssen, 2007). If this was the case, we would not observe differences in overall content, but we would observe expansions associated with a given TE group or even superfamily. However, this was not the case as we did not observe order-level expansions (Figure 2) or superfamily expansions (Figure 3). Despite the DNA/hAT superfamily being the most abundant superfamily across the radiation, there are not significant differences across species that would indicate a superfamily-specific burst following colonization to new volcanic communities. However, there are limitations to our low-coverage WGS data and analyses, as differential copy levels are not distinguishable at a finer scale with low-coverage approaches such as dnaPipeTE (Figure 4-5).

### TE composition in Tetragnatha spiny-leg radiation

The classification of TEs in Arachnid genomes has been challenging, as a high portion of unknown TEs and repeats are typically reported (e.g., Cerca et al. 2021; Wang et al 2022). This is not surprising since there are no model organisms in Arachnids (Brewer *et al*., 2014). However, spider genomes are attractive targets for those interested in TE dynamics as there is a wide variation in TE content and genome size (Garb, Sharma, & Ayoub, 2018). We found that DNA/hAT transposons are the most common type of TEs detected in the analyzed *Tetragnatha* genomes, followed by LTR/Gypsy transposons (Figure 2). This is similar to the overall repeat content of other distantly related spider lineages (Cerca *et al*., 2021; Wang *et al*., 2022). DNA/hAT transposons were the most common TE superfamily, as reconstructed by dnaPipeTE, with 2,668 contigs for the spiny leg clade on average, compared to the next most abundant superfamily, LTR/Gypsy (average of 571 elements). In the *Tetragnatha* genomes, there were 11 different hAT families, the most common being hAT-Tip100, Blackjack, Charlie, and hATm. Furthermore, despite their low numbers over *Tetragnatha* genomes, we found significant differences for the number of satellite repeats between the spiny-leg lineage to the web-builder lineage. We found that satellite elements were significantly different between spiny-legs and the web-builder clade (Table 2). This is in agreement with previous investigations that suggested that satellites could be used as phylogenetic markers, given their fast evolutionary rates (Pons & Gillespie, 2004; Roderick *et al*., 2012).

## Conclusions

The fact that species experiencing adaptive radiation on oceanic islands experience strong bottle necks, which reduce their genetic diversity, presents a paradox. What is the source of genetic variation which acts as the raw material for ecological and phenotypic diversification in oceanic archipelagos? Here, we explored whether population oscillations associated with founder events could lead to a genome shock and bursts of TEs. We did not find an overall accumulation of TEs, no specific bursts of super-families, and no differences in the age of accumulation of TEs. While we conclude that genome shock may not have facilitated an increase in the overall genetic diversity of young populations/species on the spiny-leg clade, we cannot exclude the possibility that it has acted on specific genes or specific pathways which may be associated with phenotypic and environmental diversity.

## Funding & Acknowledgements

HHY was supported by the UC Berkeley Rose Hills Independent Summer Undergraduate Research Fellowship Program. DDC was supported by a Fulbright/CONICYT Doctoral Fellowship, Integrative Biology Department and the Graduate Division of UC Berkeley, the Margaret C. Walker Fund (Essig Museum of Entomology), Sigma Xi and an Alexander von Humboldt Postdoctoral Fellowship. The authors would like to acknowledge a large number of people and institutions that collaborated at different stages of this research. The fieldwork in Hawaiʻi was supported by Laura Arnold, Timothy Bailey, David Benítez, Katie Champlin, James Friday, Jun Ying Lim, Emory Griffin-Noyes, Faith Inman-Narahari, Darcey Iwashita, Raina Kaholoaa, Jessie Knowlton, Rick Lapoint, Scott Laursen, Karl Magnacca, Elizabeth Morrill, Patrick O’Grady, Rita Pregana, Donald Price, David Rankin, William Roderick, Karen Uy, Erin Wilson-Rankin and the Kīpuka team.

The permit processing and access to different reserves and private land was possible thanks to Steve Bergfeld (DOFAW Big Island), Pat Bily (TNC Maui), Tabetha Block (HETF), Shalan Crysdale (TNC Big Island), Lance DaSilva (DOFAW Maui), Danae Dean (Kahoma Ranch), Charmian Dang (NAR), Melissa Dean (HETF), Betsy Gagne (NAR), Elizabeth Gordon (HALE), Lisa Hadway (DOFAW Big Island), Paula Hartzell (Lanaʻi Resorts, LLC), Greg Hendrickson (Kealakekua Ranch), Mel Johansen (TNC Big Island), Pomaikaʻi Kaniaupio-Crozier (Maui Land and Pinneapple), Cynthia King (DLNR), Peter Landon (NAR Maui), Rhonda Loh (HAVO), Russell Kallstrom (TNC Molokaʻi), Joey Mello (DOFAW Big Island), Ed Misaki (TNC Molokaʻi), Elliot Parsons (Puʻu Waʻawaʻa HETF), Lani Petrie (Kapapala Ranch), Shawn Saito (Parker Ranch), Joe Ward (Maui Land and Pinneapple) and Kawika Winter (Limahuli Botanical Garden).

## Data Availability Statement

The data used in this article are available in the Genbank BioProject database at https://www.ncbi.nlm.nih.gov/bioproject/PRJEB64196/, and can be accessed with project accession number PRJEB64196.

## Appendix

**Supplementary Table 01.** Sampling information and public (ENA) accession IDs. (EM – East Maui; Kil – Kilauea; ML – Mauna Loa; Koh – Kohala; Mo – Moloka’i; Wai – Wai’anae; Ka – Kaua’i; Koo – Ko’olau)

| Species | Island | Volcano | ID | study_accession | sample_accession | experiment_accession | run_accession | tax_id |
| --- | --- | --- | --- | --- | --- | --- | --- | --- |
| *T. acuta* | Maui | EM | 071 | PRJEB64196 | SAMEA114006635 | ERX11079388 | ERR11678375 | 172563 |
| *T. anuenue* | Hawaii | Kil | 048 | PRJEB64196 | SAMEA114006638 | ERX11079389 | ERR11678376 | 263990 |
| *T. anuenue* | Hawaii | ML | 066 | PRJEB64196 | SAMEA114006641 | ERX11079517 | ERR11678504 | 263990 |
| *T. brevignatha* | Hawaii | Kil | 007 | PRJEB64196 | SAMEA114006642 | ERX11079550 | ERR11678545 | 172555 |
| *T. brevignatha* | Hawaii | Koh | 050 | PRJEB64196 | SAMEA114006643 | ERX11079551 | ERR11678546 | 172555 |
| *T. brevignatha* | Hawaii | ML | 067 | PRJEB64196 | SAMEA114006645 | ERX11079801 | ERR11678800 | 172555 |
| *T. brevignatha* | Maui | EM | 059 | PRJEB64196 | SAMEA114006644 | ERX11079674 | ERR11678673 | 172555 |
| *T. filiciphila* | Maui | EM | 074 | PRJEB64196 | SAMEA114006646 | ERX11079841 | ERR11678840 | 300521 |
| *T. kamakou* | Maui | EM | 012 | PRJEB64196 | SAMEA114006651 | ERX11079802 | ERR11678801 | 172554 |
| *T. kamakou* | Molokai | Mo | 010 | PRJEB64196 | SAMEA114006649 | ERX11079780 | ERR11678779 | 172554 |
| *T. kauaiensis* | Kauai | Ka | 020 | PRJEB64196 | SAMEA114006659 | ERX11080263 | ERR11679262 | 156853 |
| *T. kikokiko* | Maui | EM | 072 | PRJEB64196 | SAMEA114006660 | ERX11080276 | ERR11679275 | 264991 |
| *T. kukuiki* | Oahu | Wai | 058 | PRJEB64196 | SAMEA114006662 | ERX11080065 | ERR11679064 | 264993 |
| *T. maka* | Kauai | Ka | 052 | PRJEB64196 | SAMEA114006664 | ERX11080507 | ERR11679506 | 302253<br>2 |
| *T. mohihi* | Kauai | Ka | 022 | PRJEB64196 | SAMEA114006666 | ERX11080510 | ERR11679509 | 302253<br>2 |
| *T. obscura* | Hawaii | ML | 006 | PRJEB64196 | SAMEA114006668 | ERX11080552 | ERR11679551 | 264994 |
| *T. perreirai* | Oahu | Wai | 024 | PRJEB64196 | SAMEA114006669 | ERX11080536 | ERR11679535 | 172557 |
| *T. pilosa* | Kauai | Ka | 028 | PRJEB64196 | SAMEA114006672 | ERX11080511 | ERR11679510 | 172561 |
| *T. quasimodo* | Hawaii | Koh | 004 | PRJEB64196 | SAMEA114006680 | ERX11082030 | ERR11681029 | 172560 |
| *T. quasimodo* | Hawaii | ML | 005 | PRJEB64196 | SAMEA114006681 | ERX11081963 | ERR11680962 | 172560 |
| *T. quasimodo* | Maui | EM | 064 | PRJEB64196 | SAMEA114006692 | ERX11082219 | ERR11681218 | 172560 |
| *T. quasimodo* | Molokai | Mo | 002 | PRJEB64196 | SAMEA114006678 | ERX11081936 | ERR11680935 | 172560 |
| *T. quasimodo* | Oahu | Wai | 033 | PRJEB64196 | SAMEA114006683 | ERX11081956 | ERR11680955 | 172560 |
| *T. restricta* | Maui | EM | 037 | PRJEB64196 | SAMEA114006695 | ERX11082808 | ERR11681863 |  |
| *T. stellarobusta* | Maui | EM | 001 | PRJEB64196 | SAMEA114006694 | ERX11082881 | ERR11681936 | 300528 |
| *T. tantalus* | Oahu | Koo | 039 | PRJEB64196 | SAMEA114006699 | ERX11082759 | ERR11681814 | 172559 |
| *T. waikamoi* | Maui | EM | 062 | PRJEB64196 | SAMEA114006706 | ERX11082831 | ERR11681886 | 35633 |

**Supplementary Table 02.**
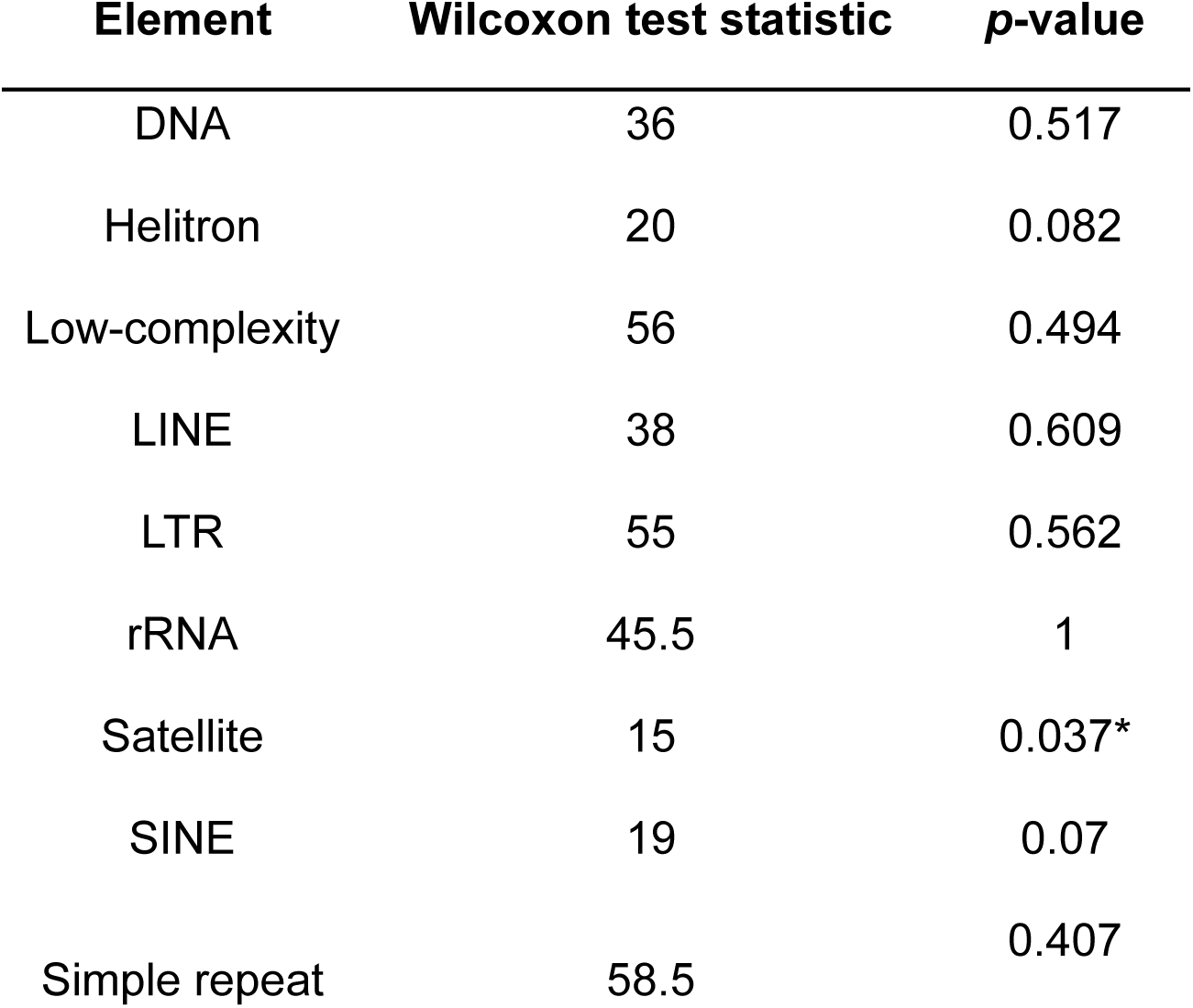
Statistical outcomes of Mann-Whitney U-test comparing abundance of repeat element groups between spiny-leg and web-builder individuals.

**Supplementary Figure 1.**
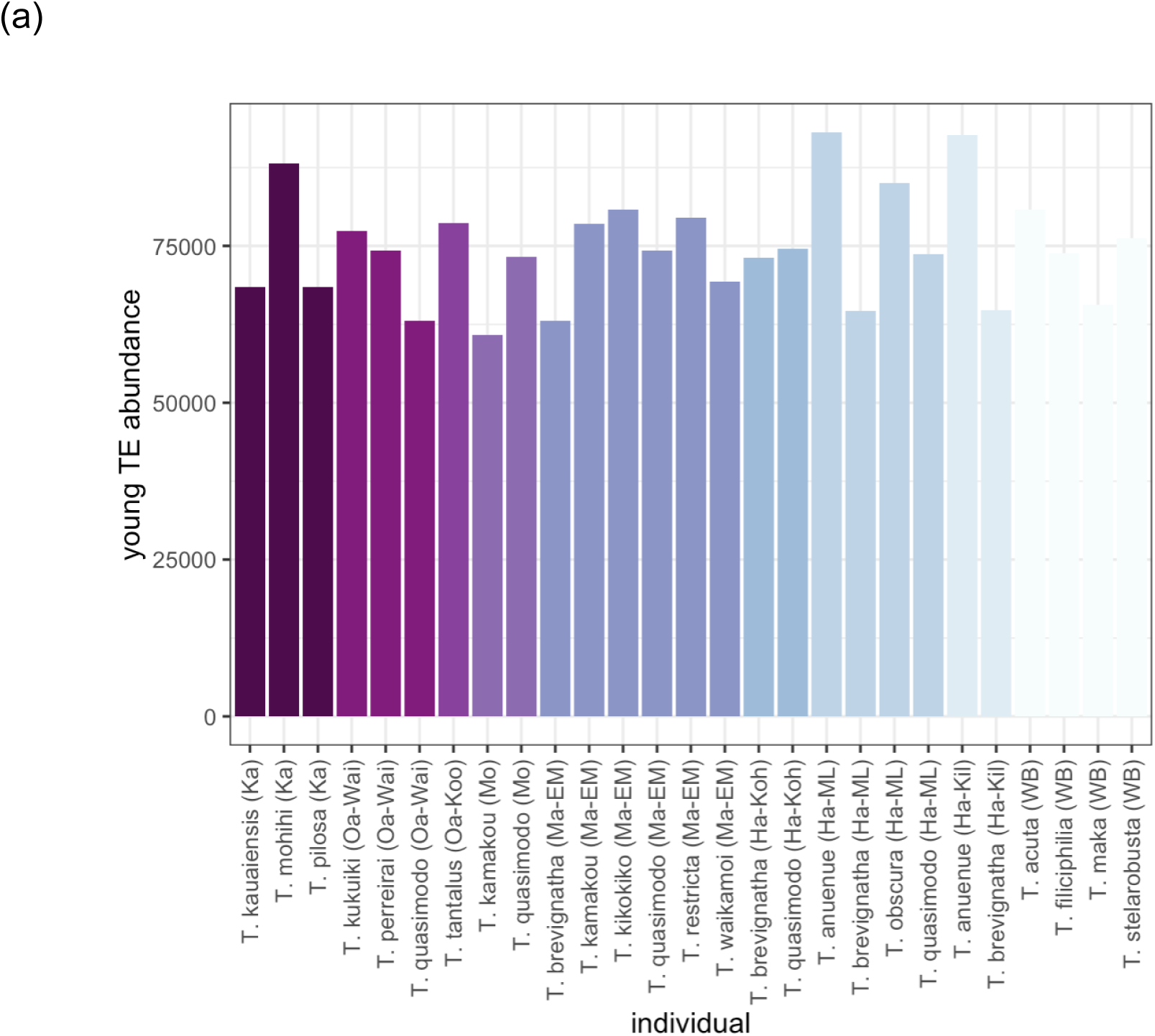
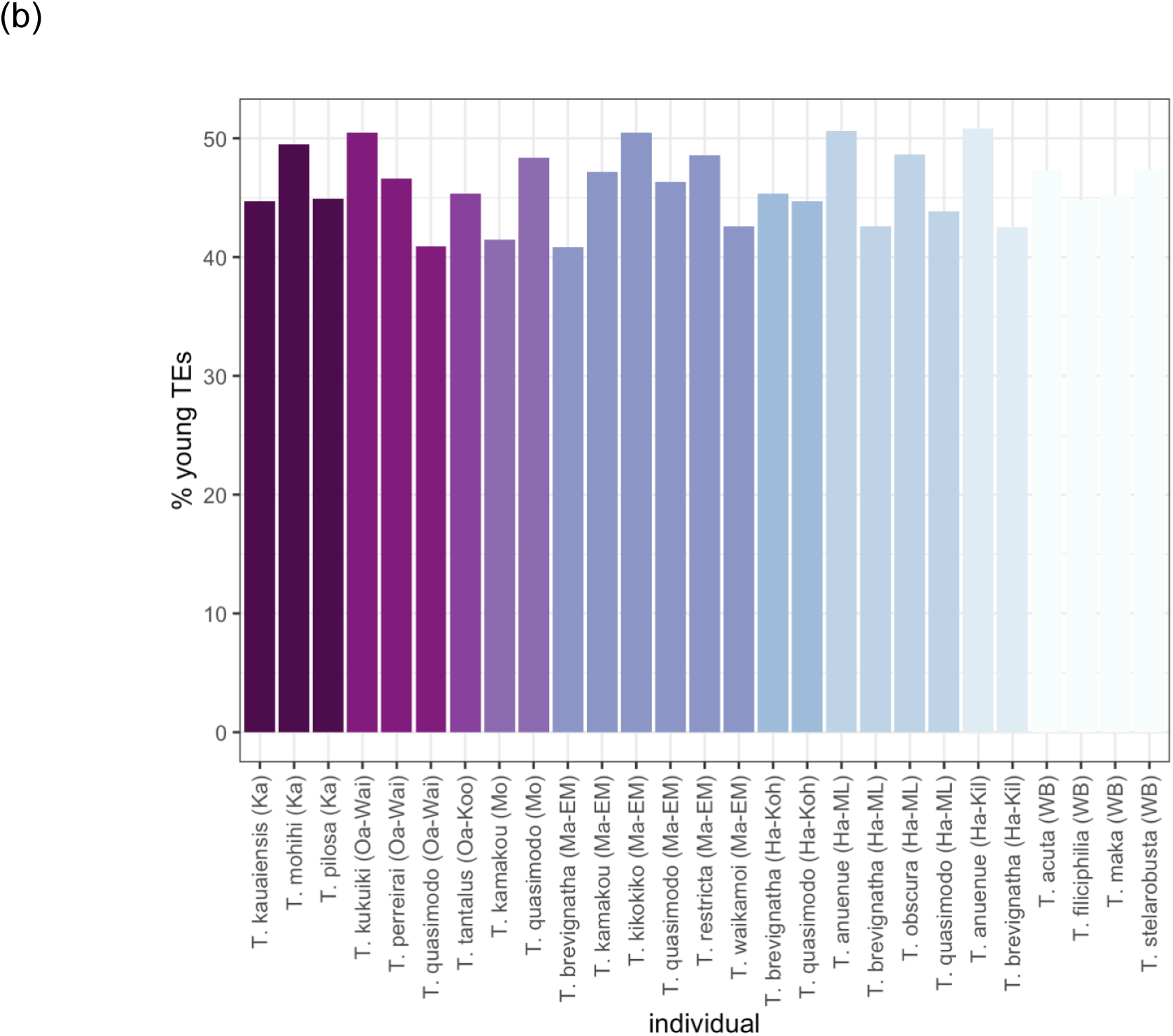
(a) Abundance and (b) percentage of young TEs within all *Tetragnatha* individuals. Similar to the percentage of TEs <5% divergent from their consensus sequences, there is no increasing trend of young TE abundance with decreasing volcanic age. *T. anuenue* of Kilauea and Mauna Loa has the highest abundance, while *T. kamakou* on Molokai has the lowest abundance.

**Supplementary Figure 2.**
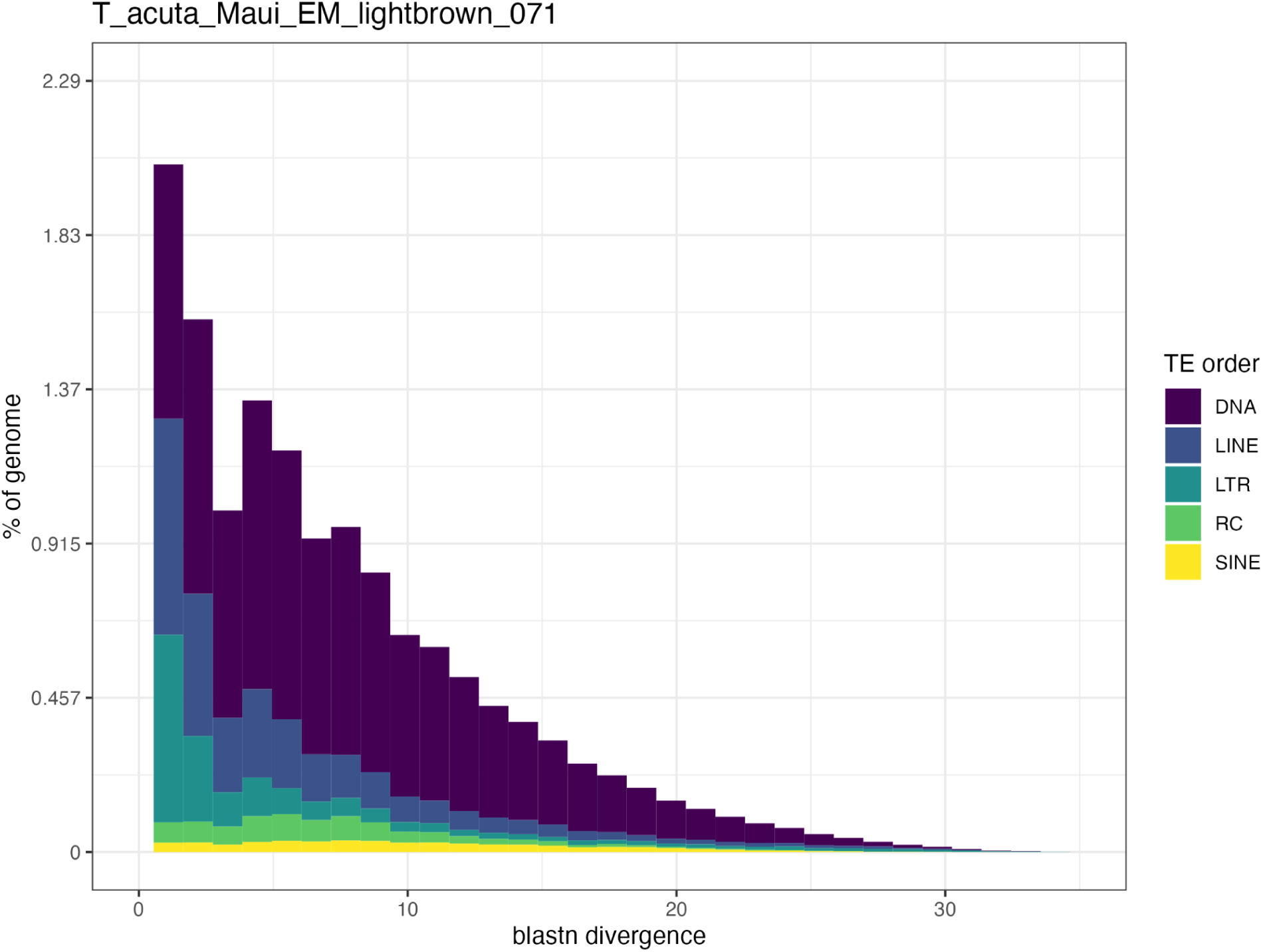
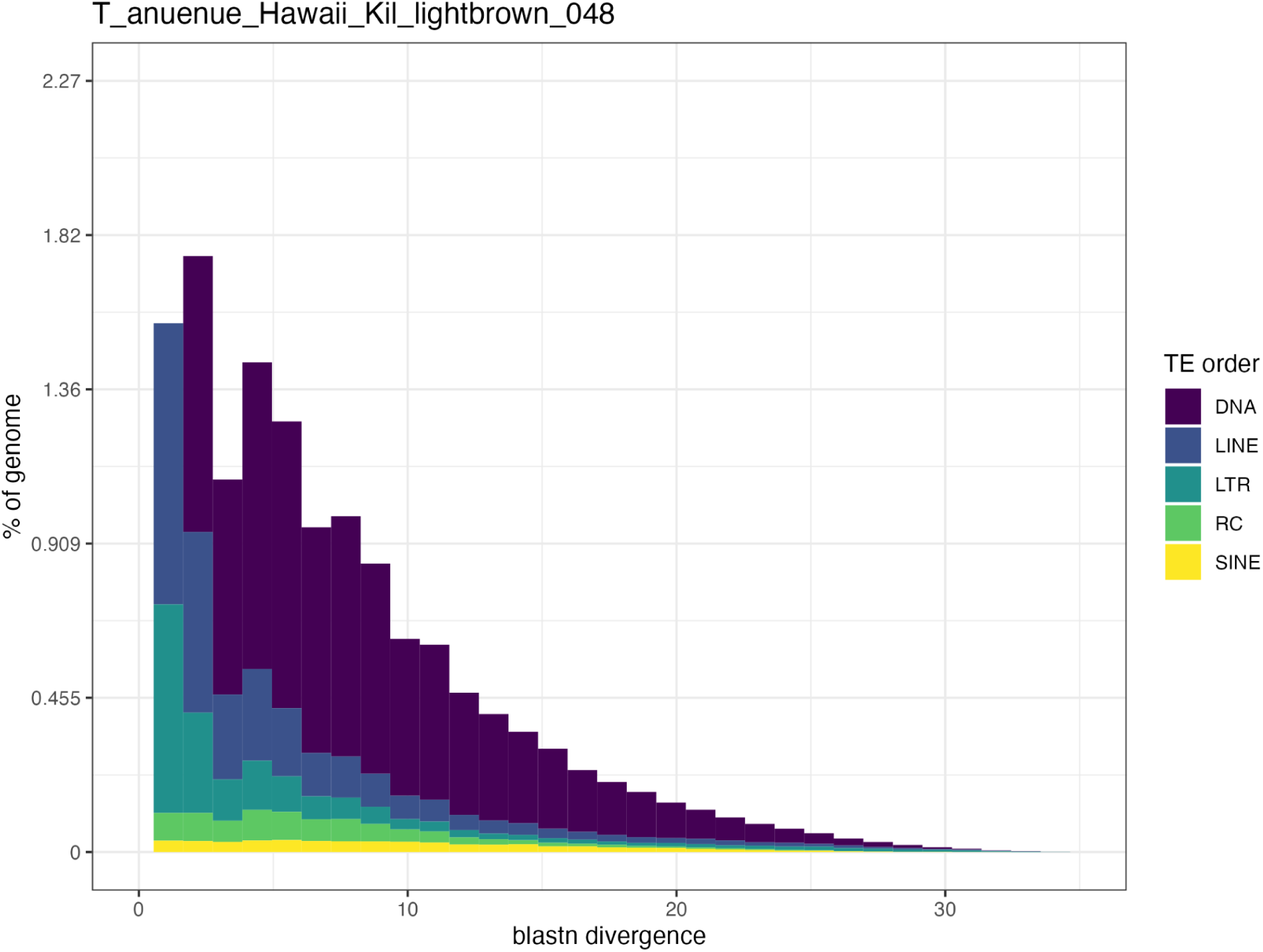
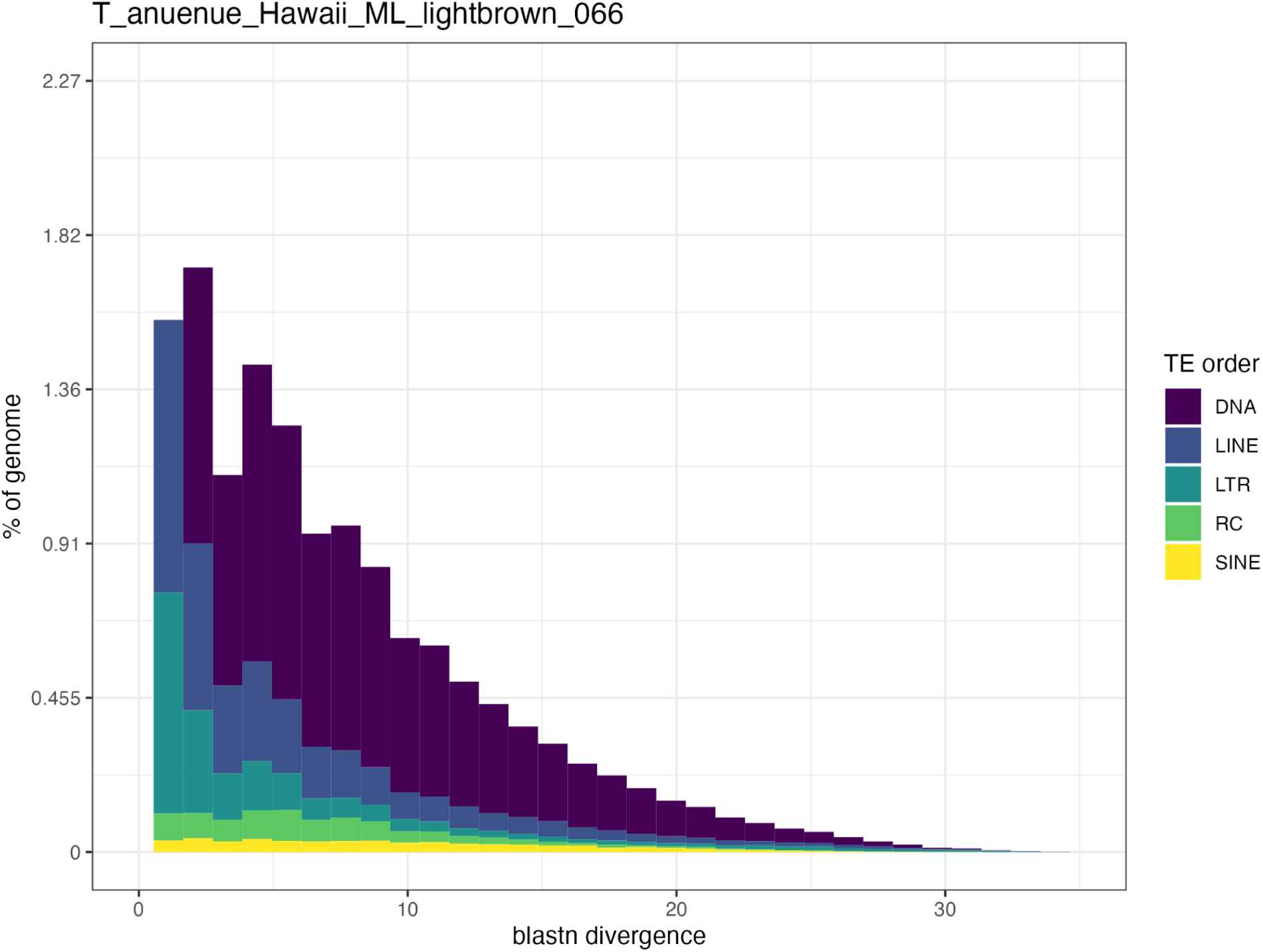
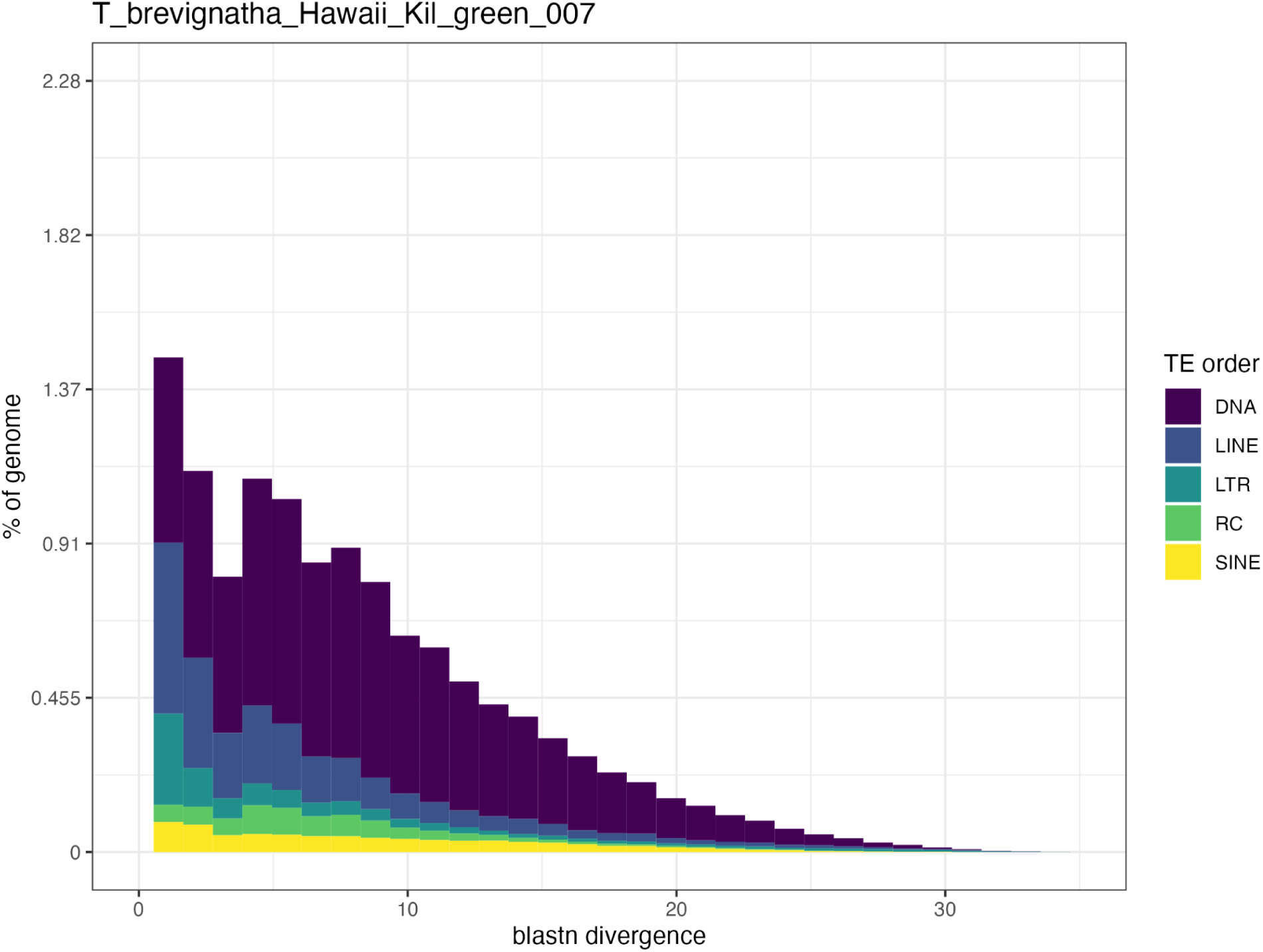
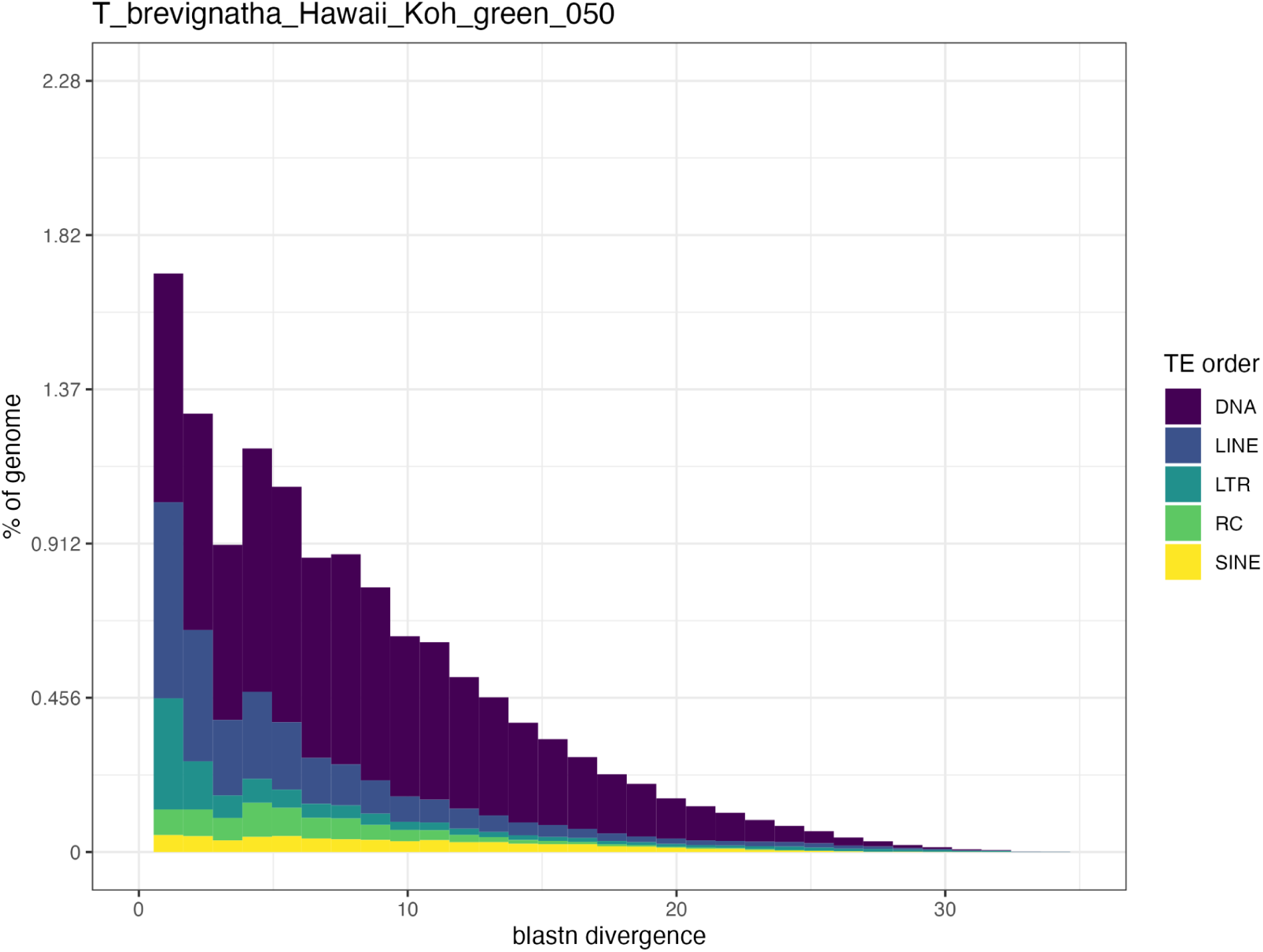
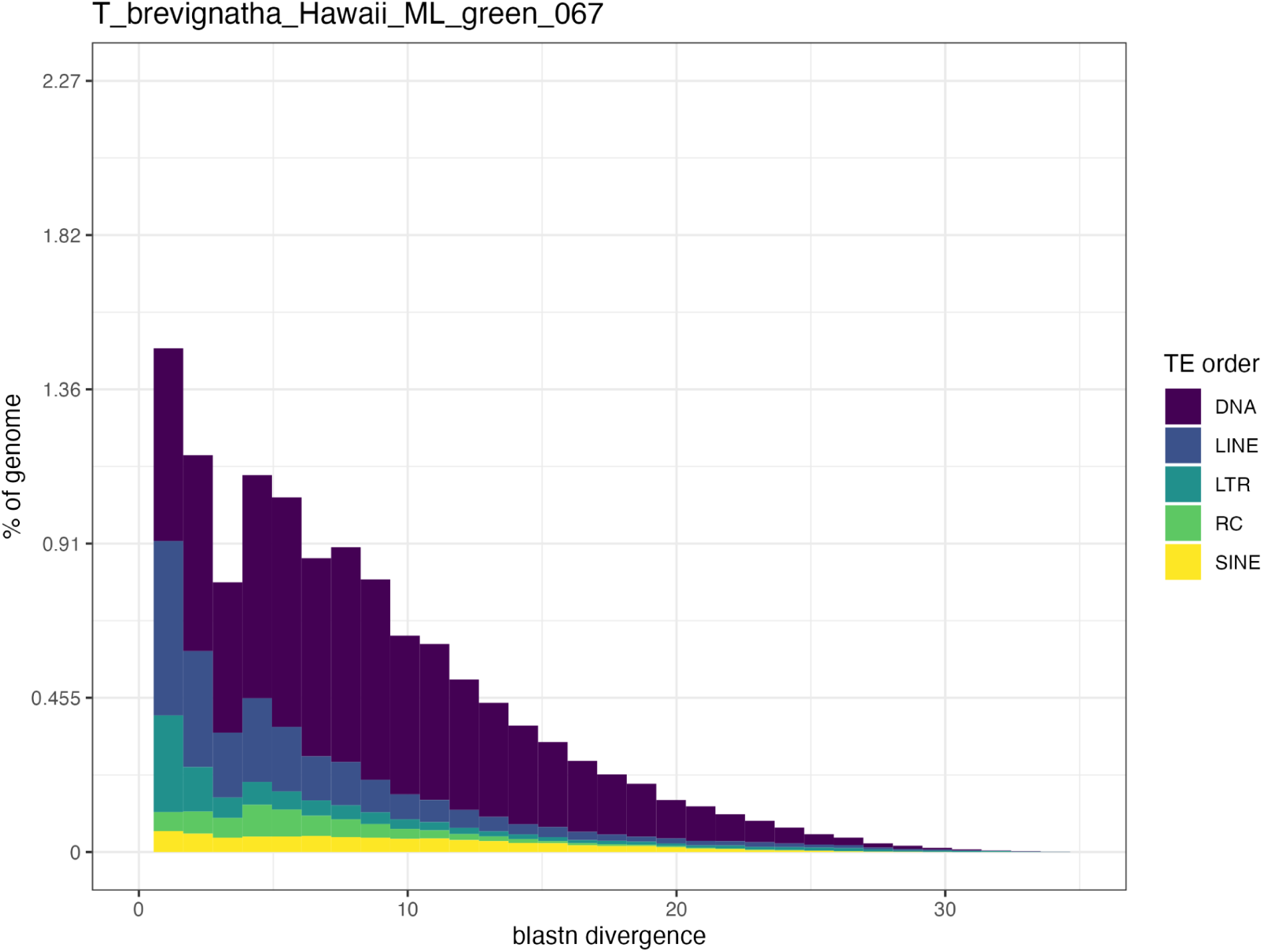
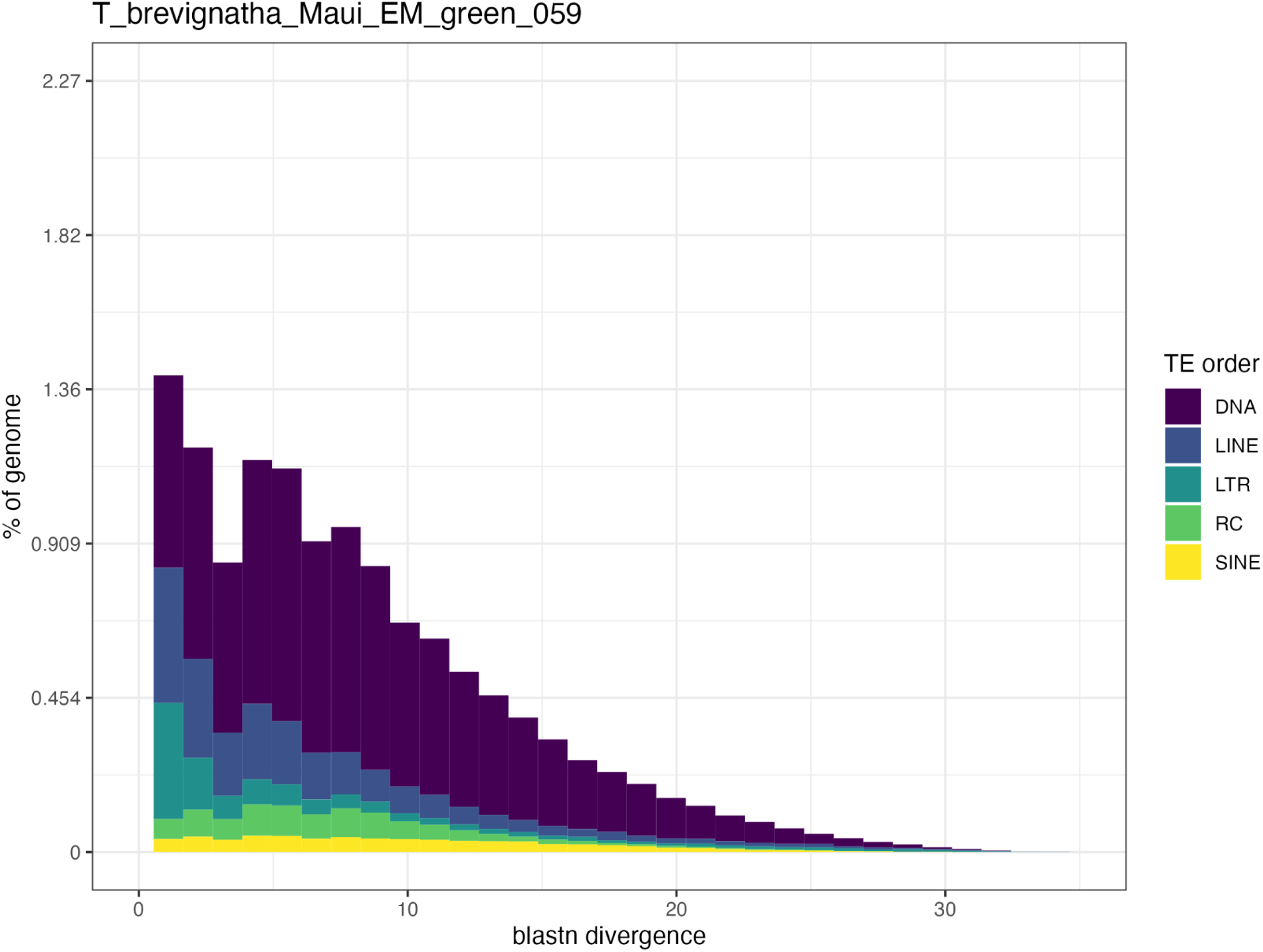
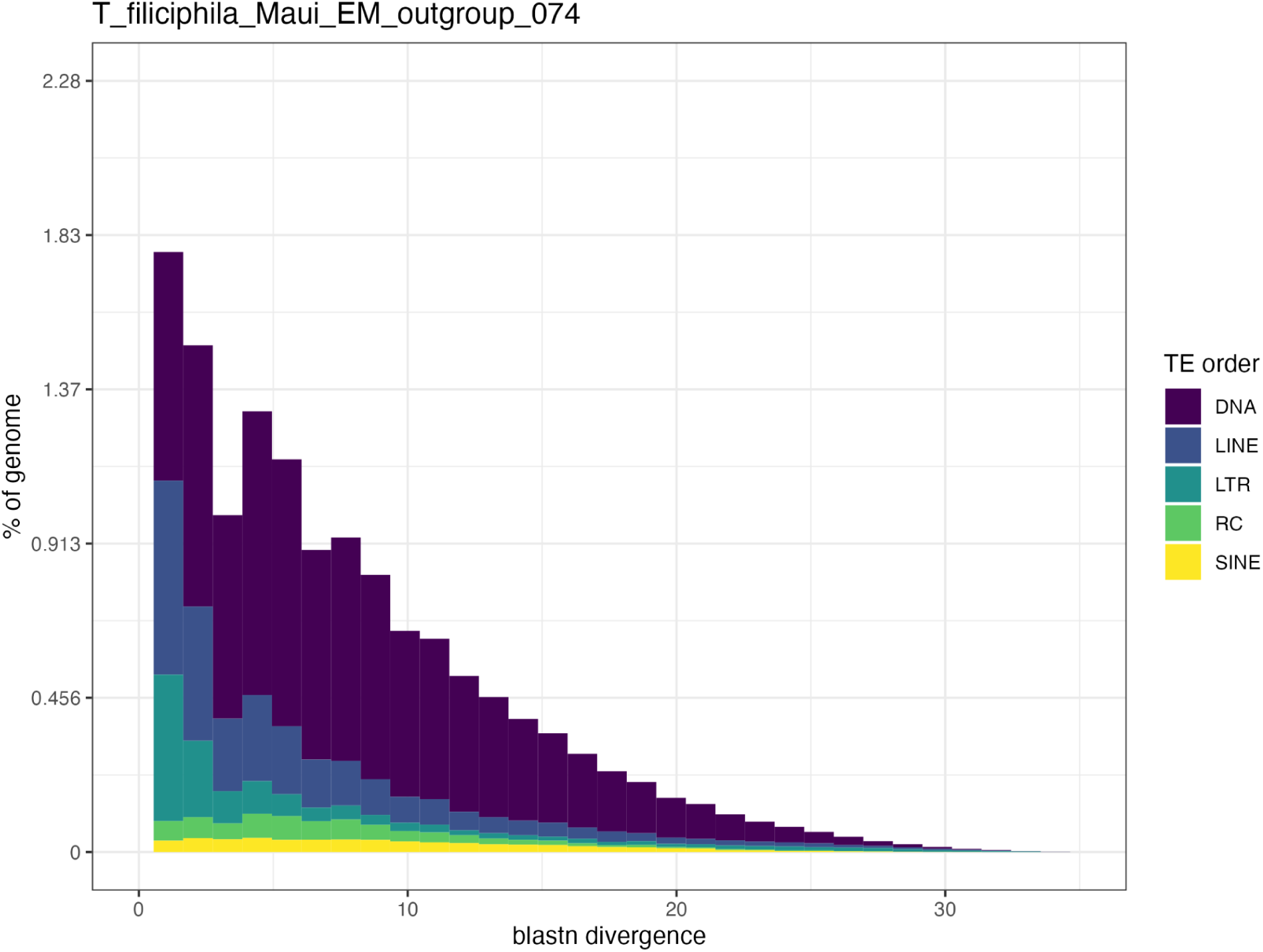
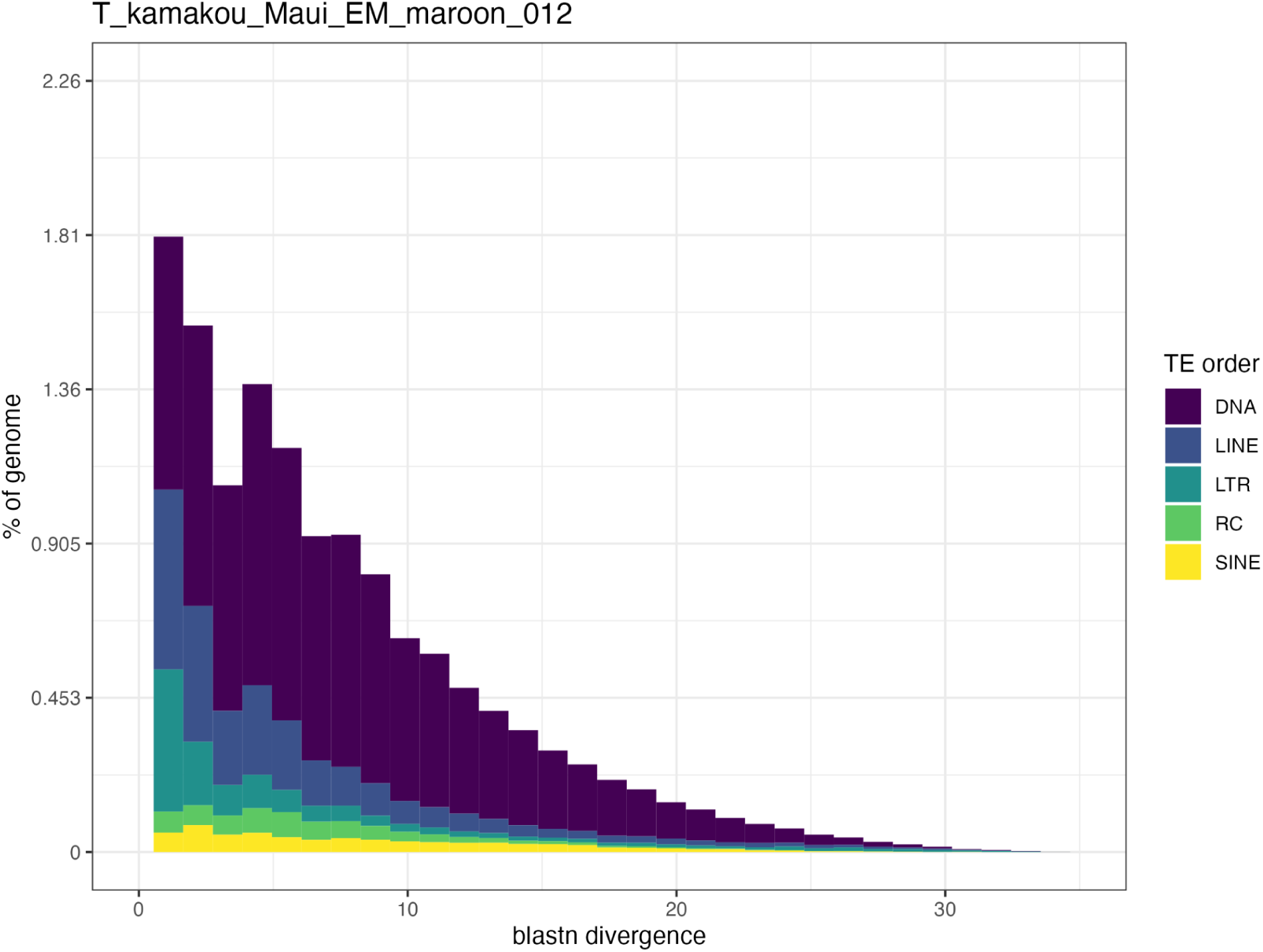
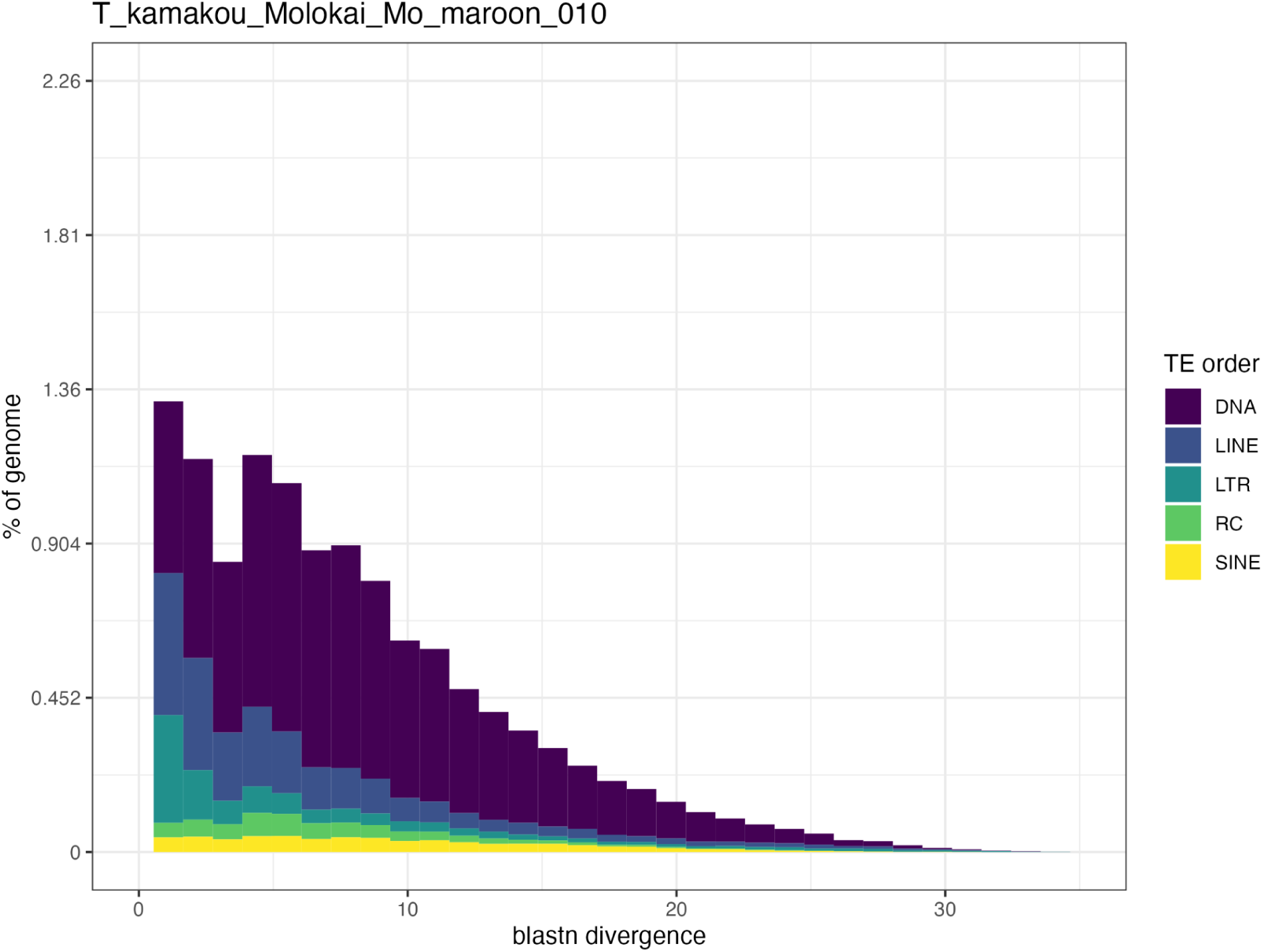
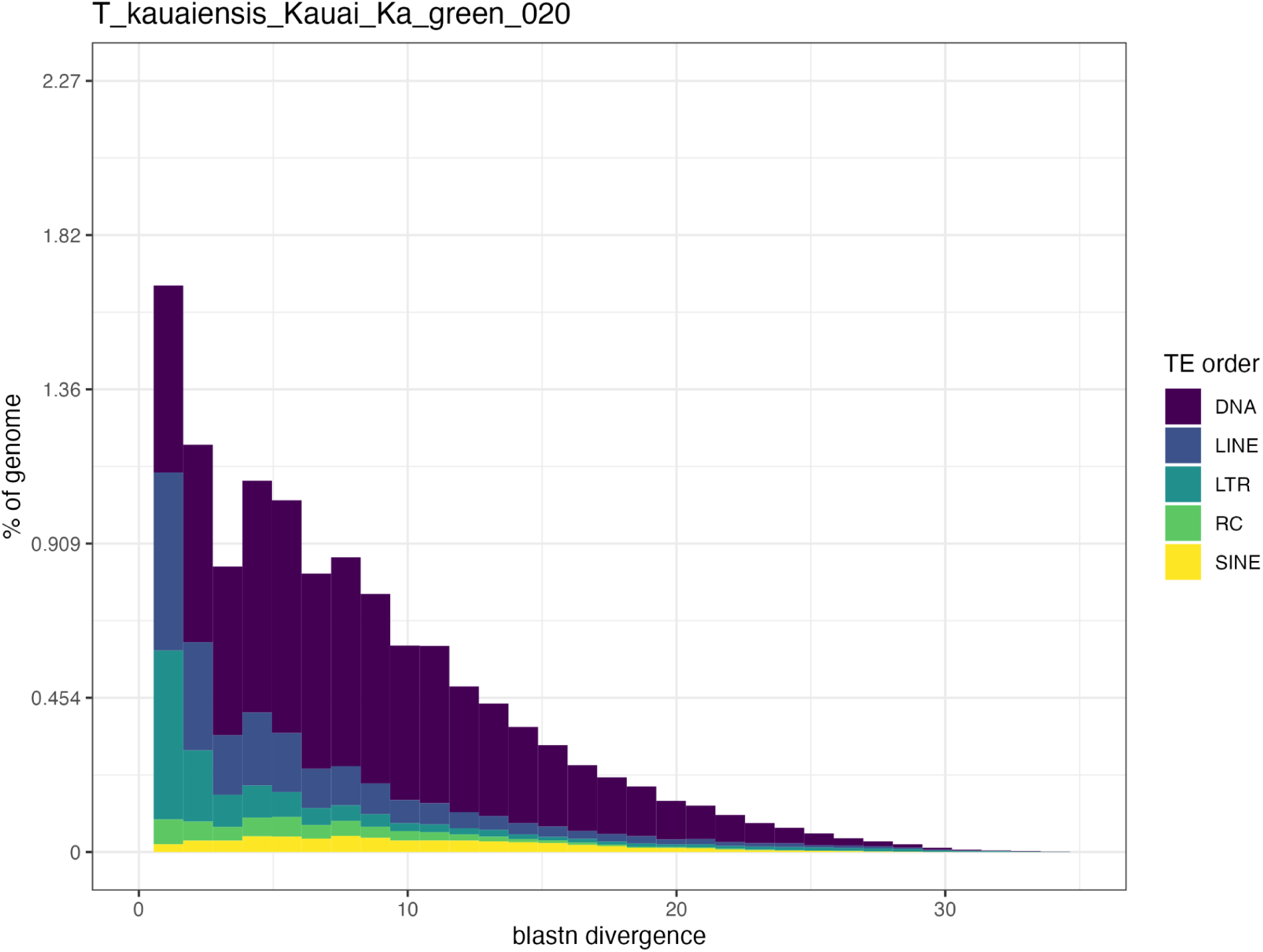
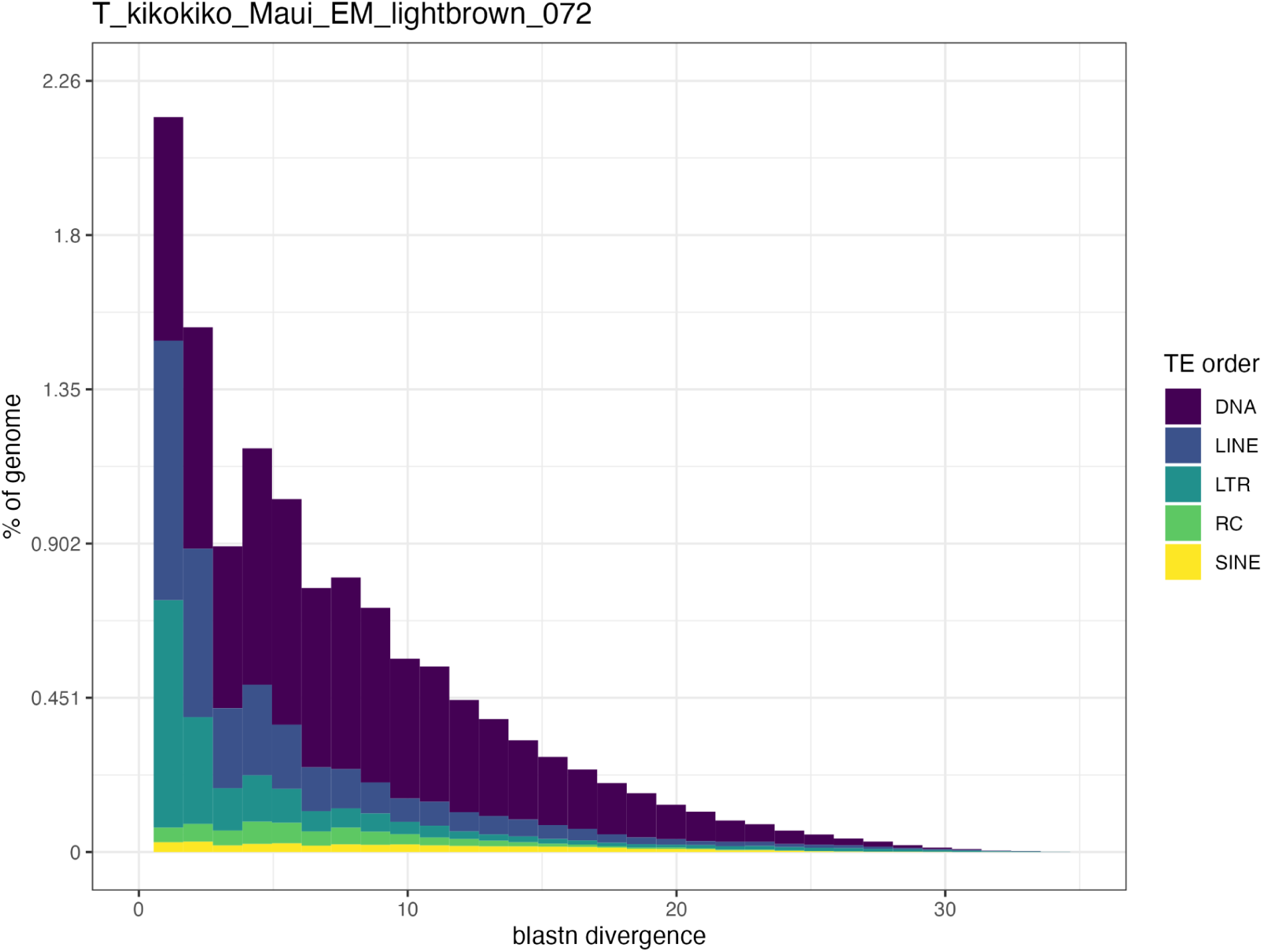
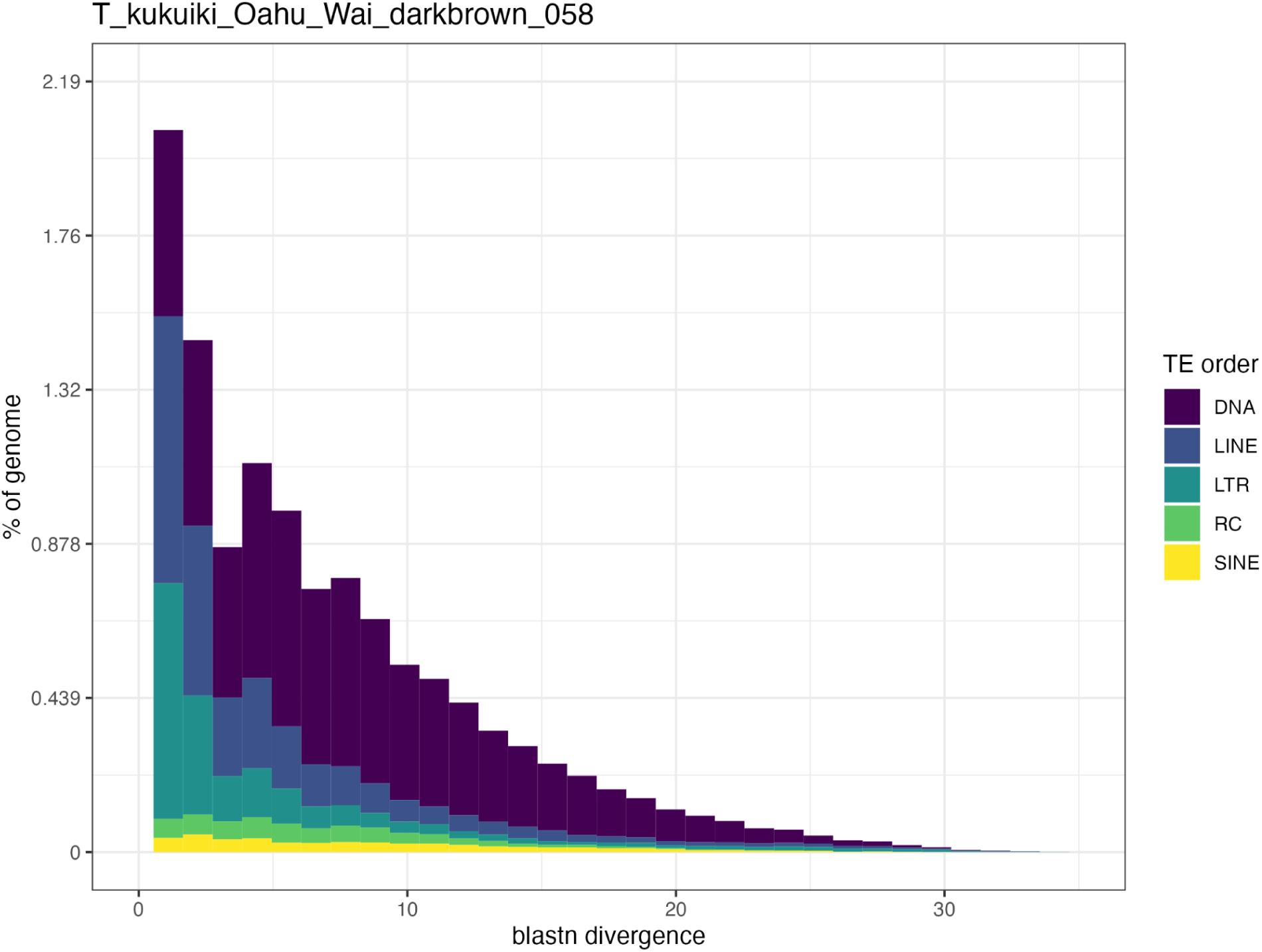
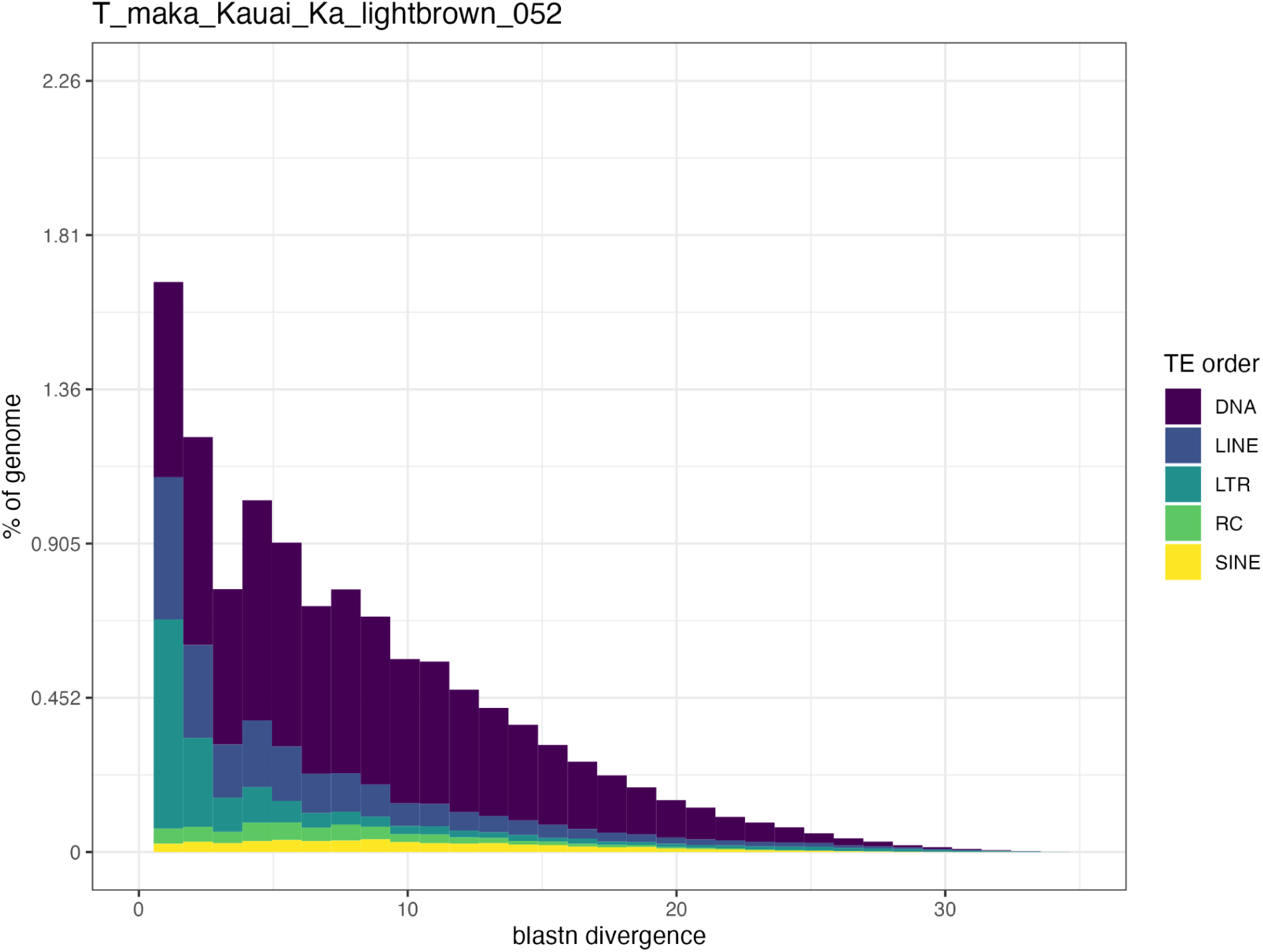
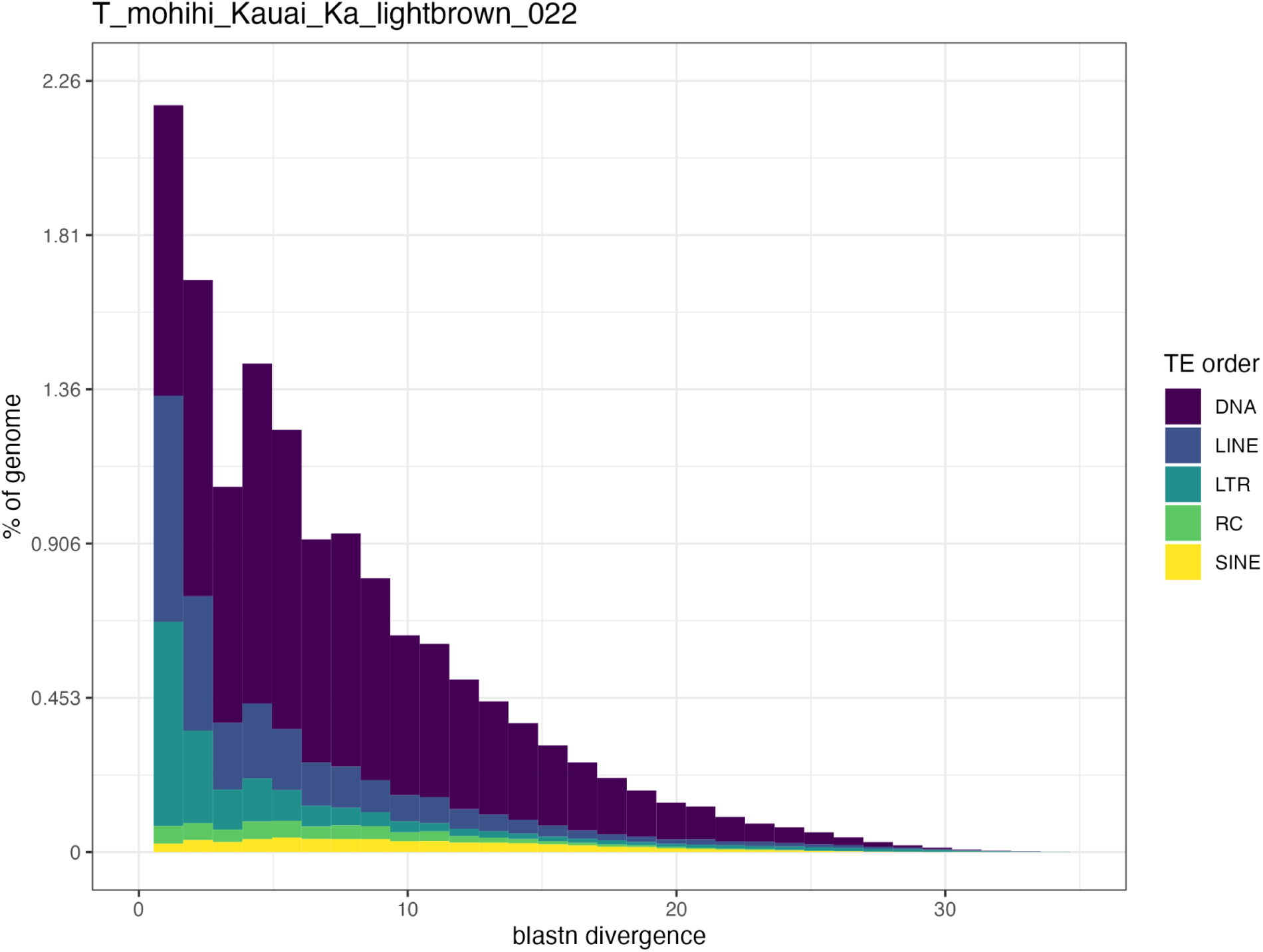
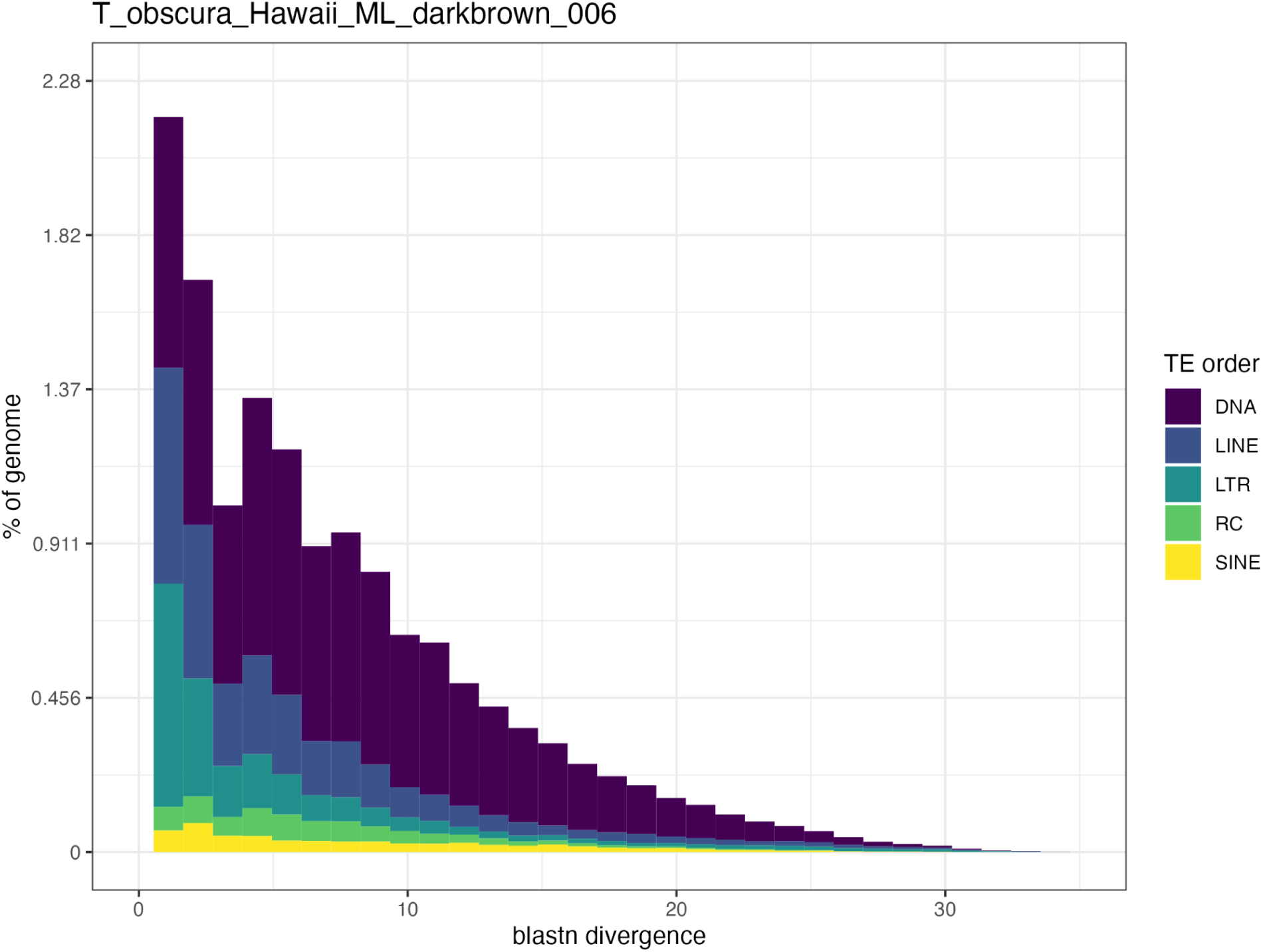
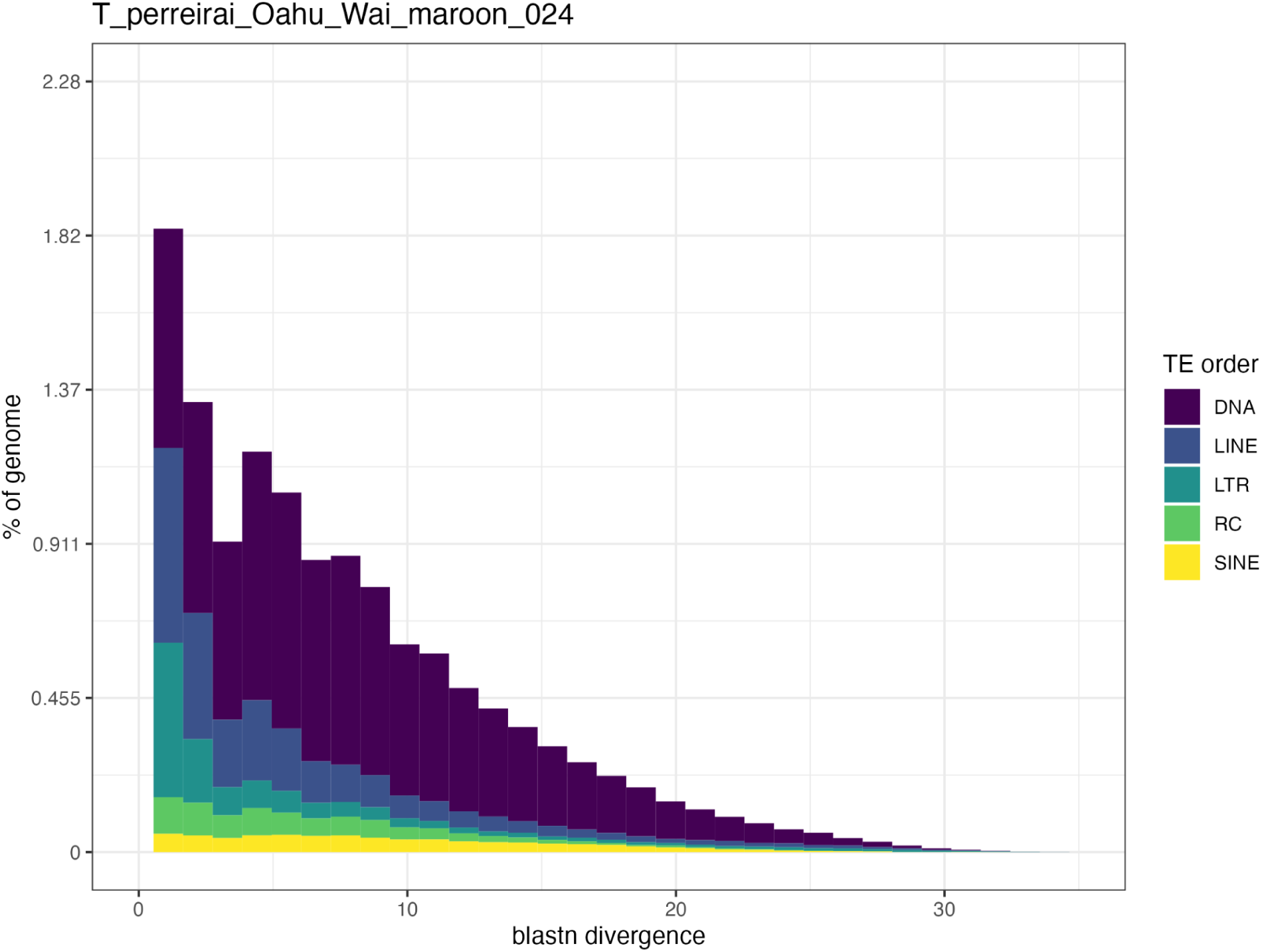
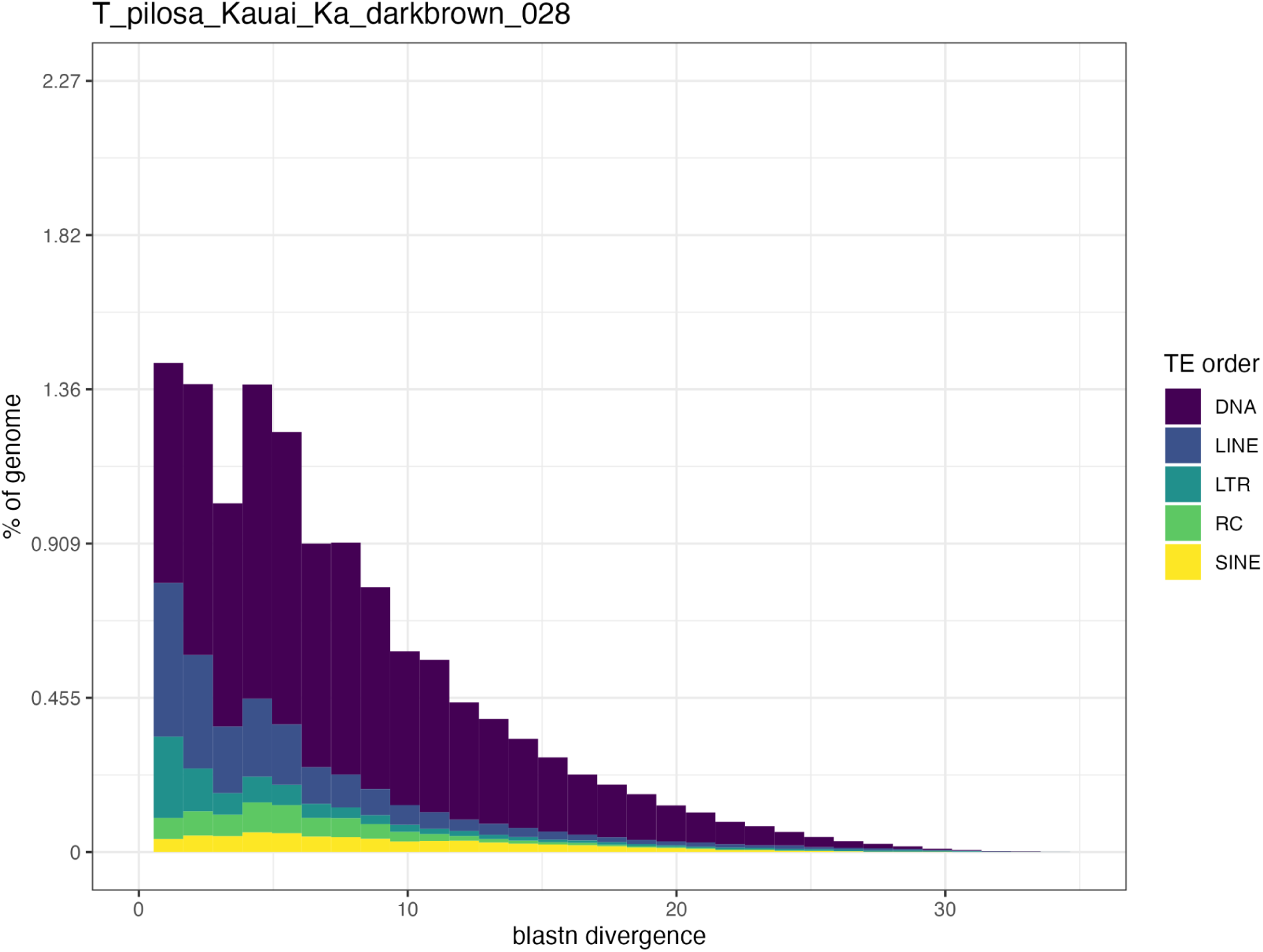
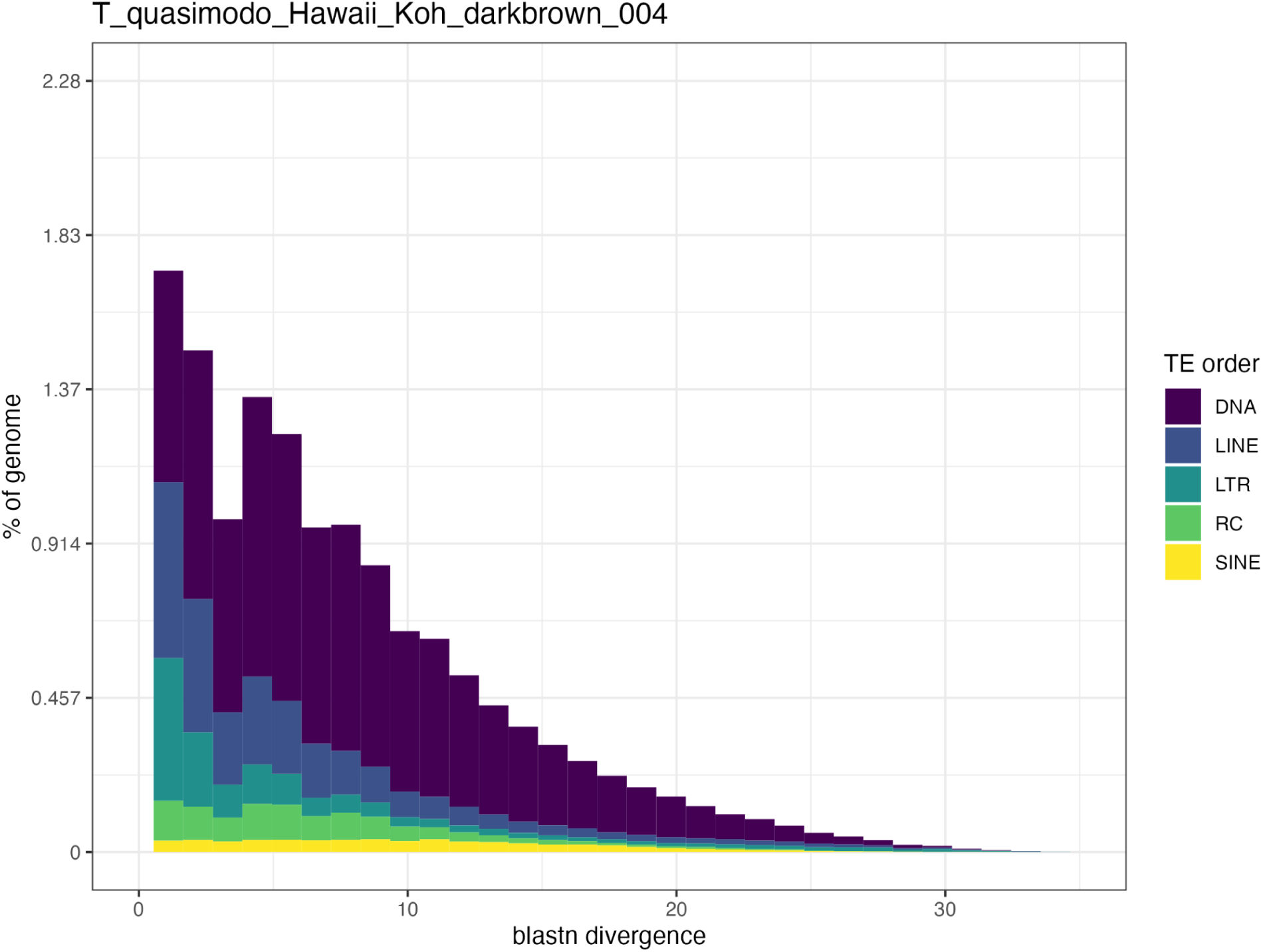
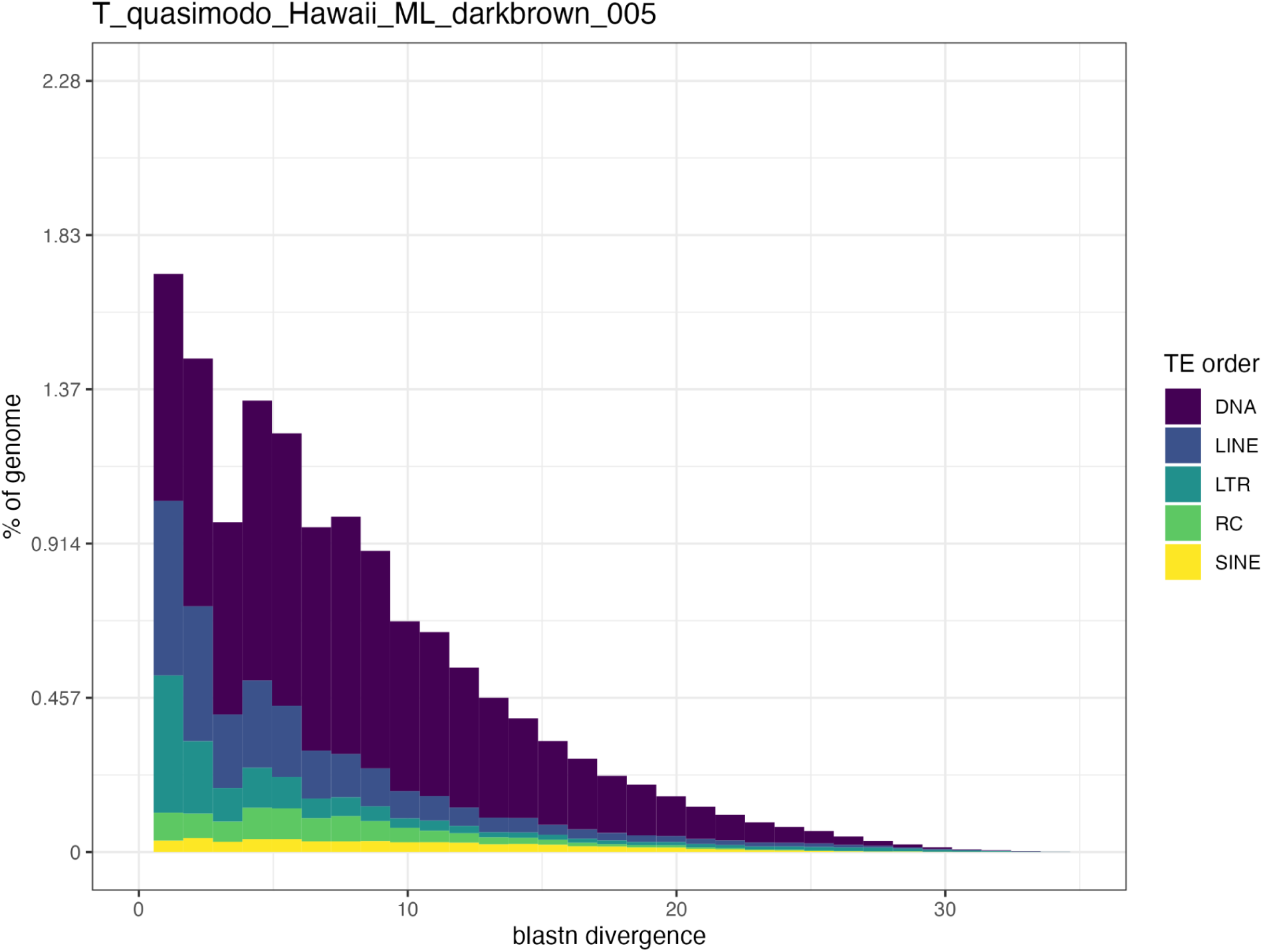
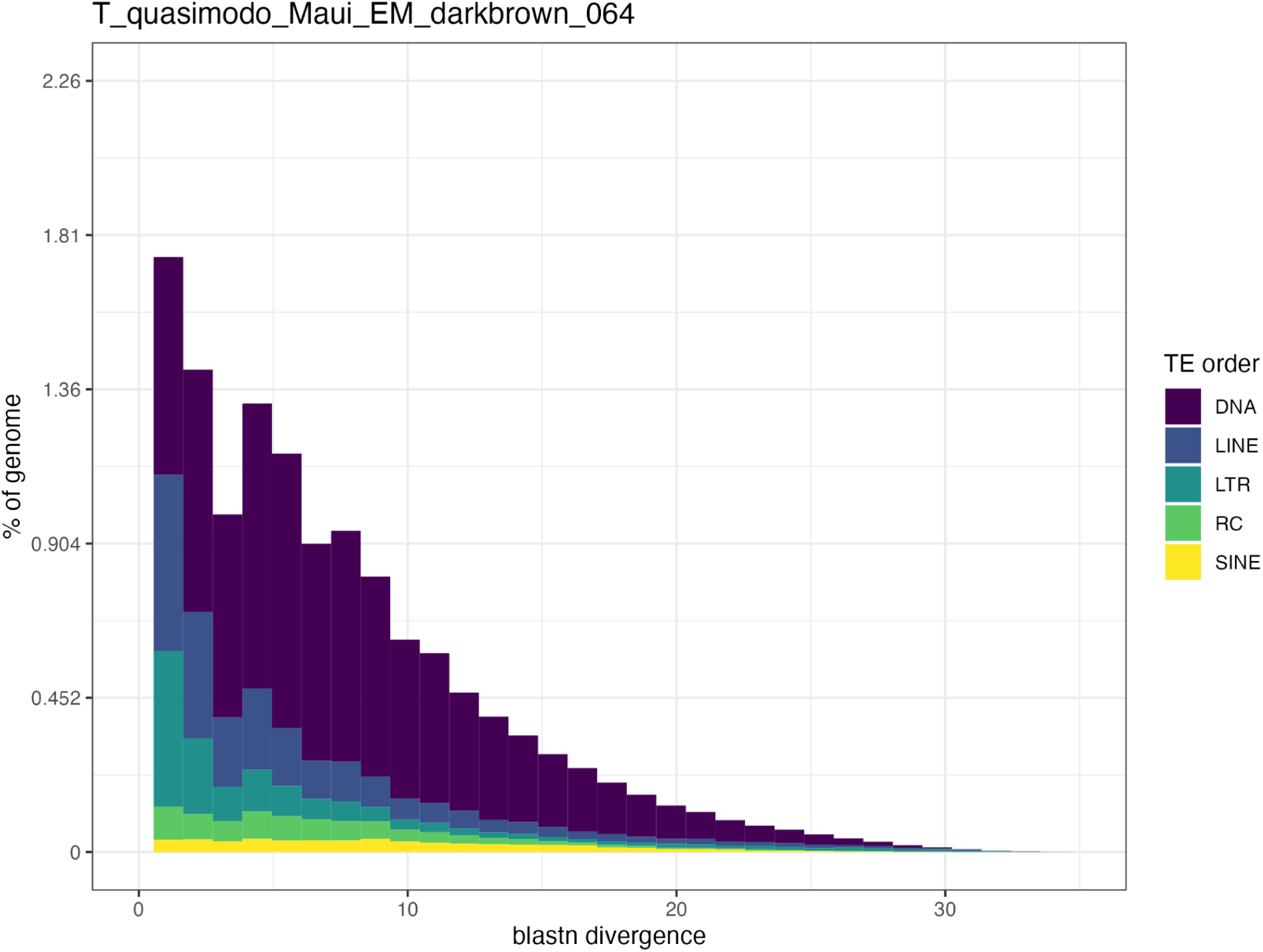
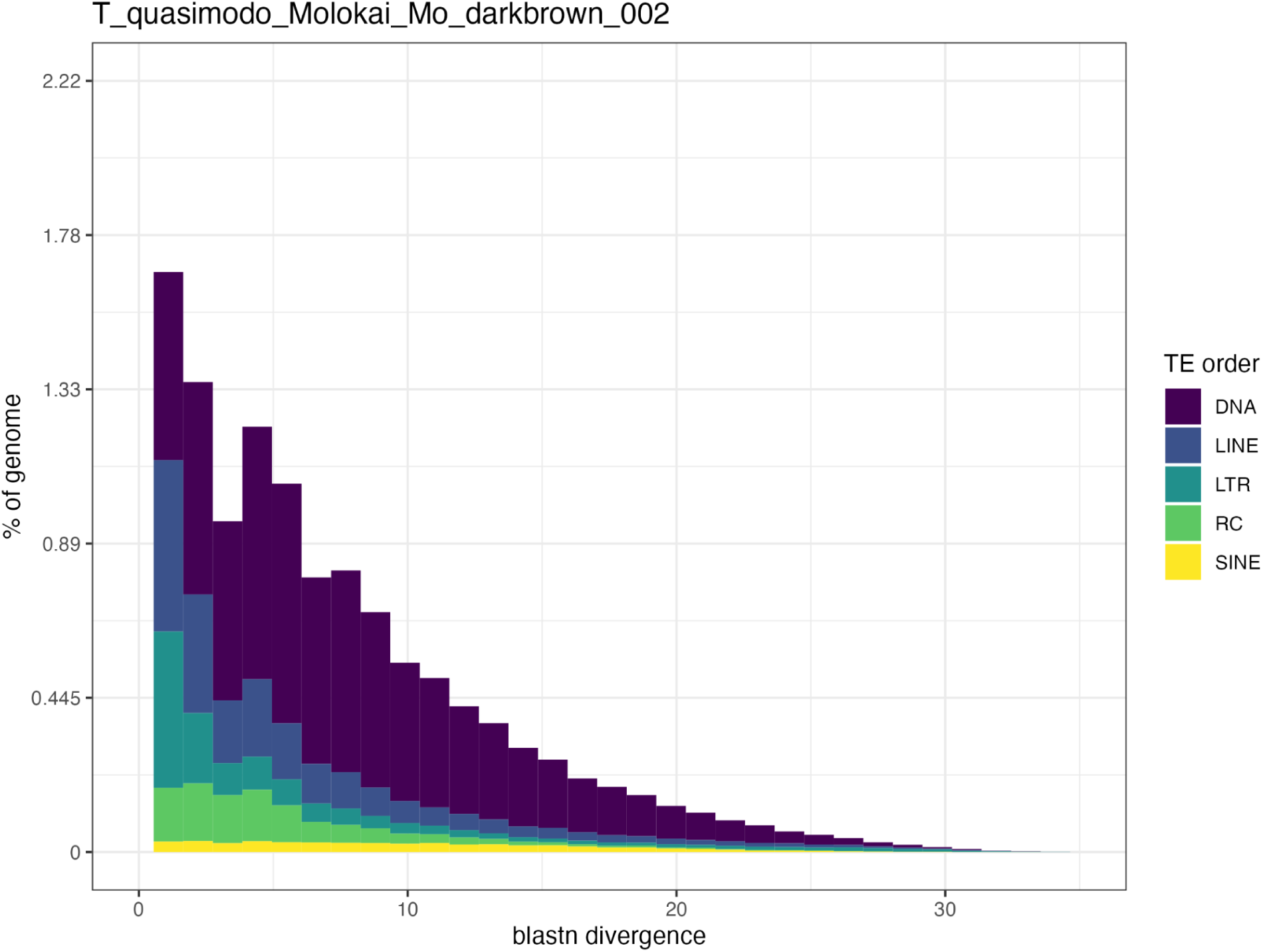
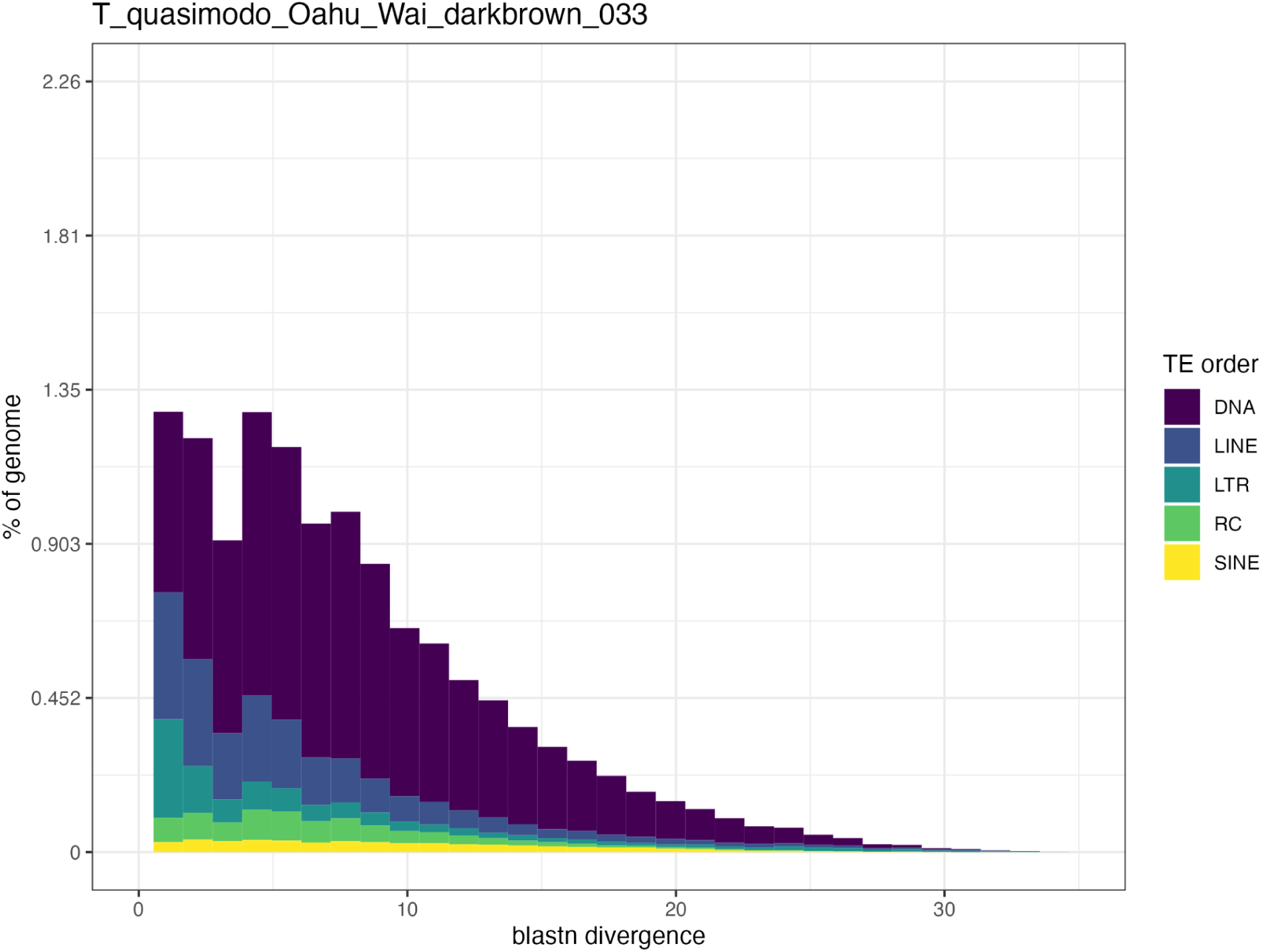
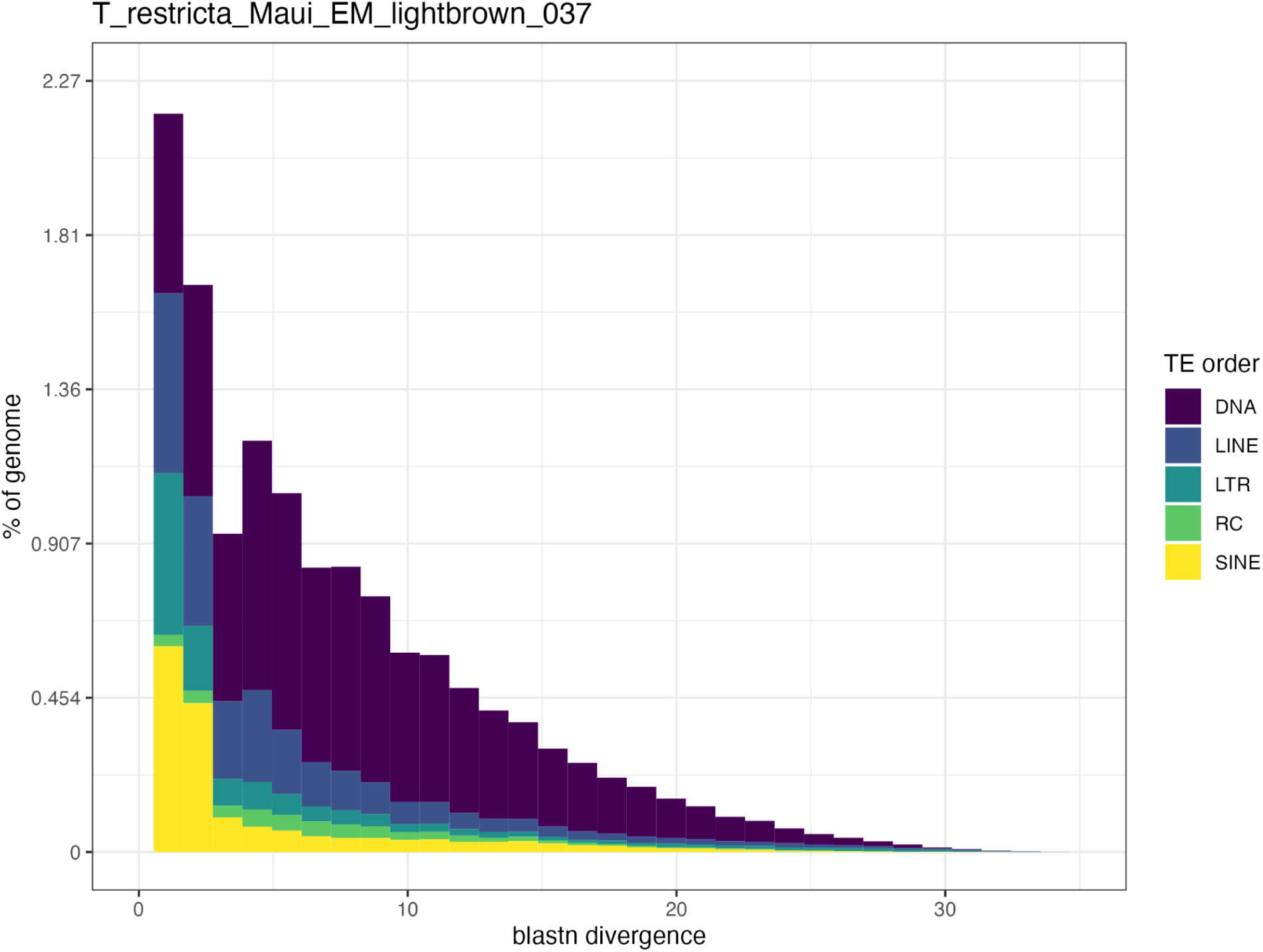
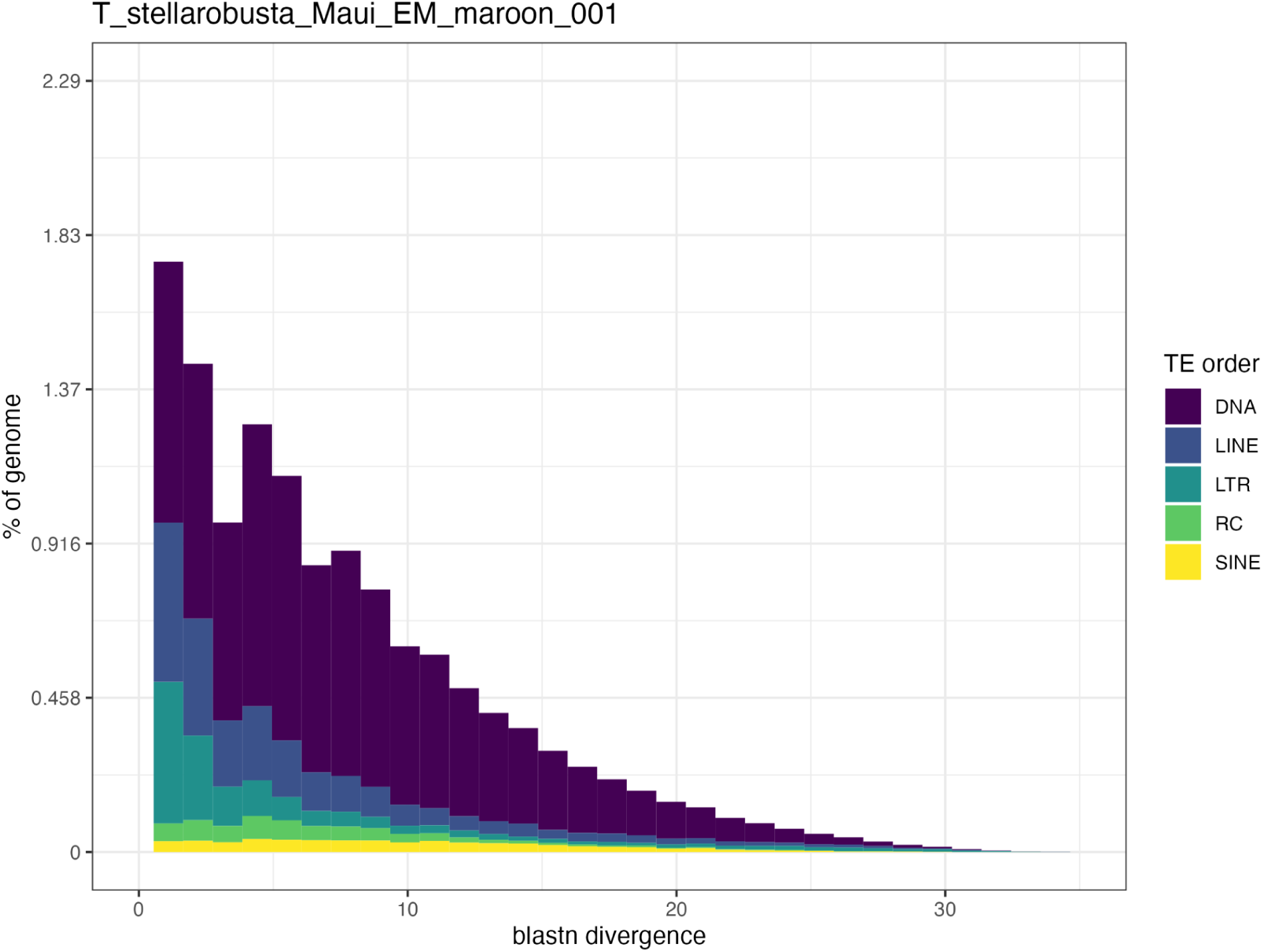
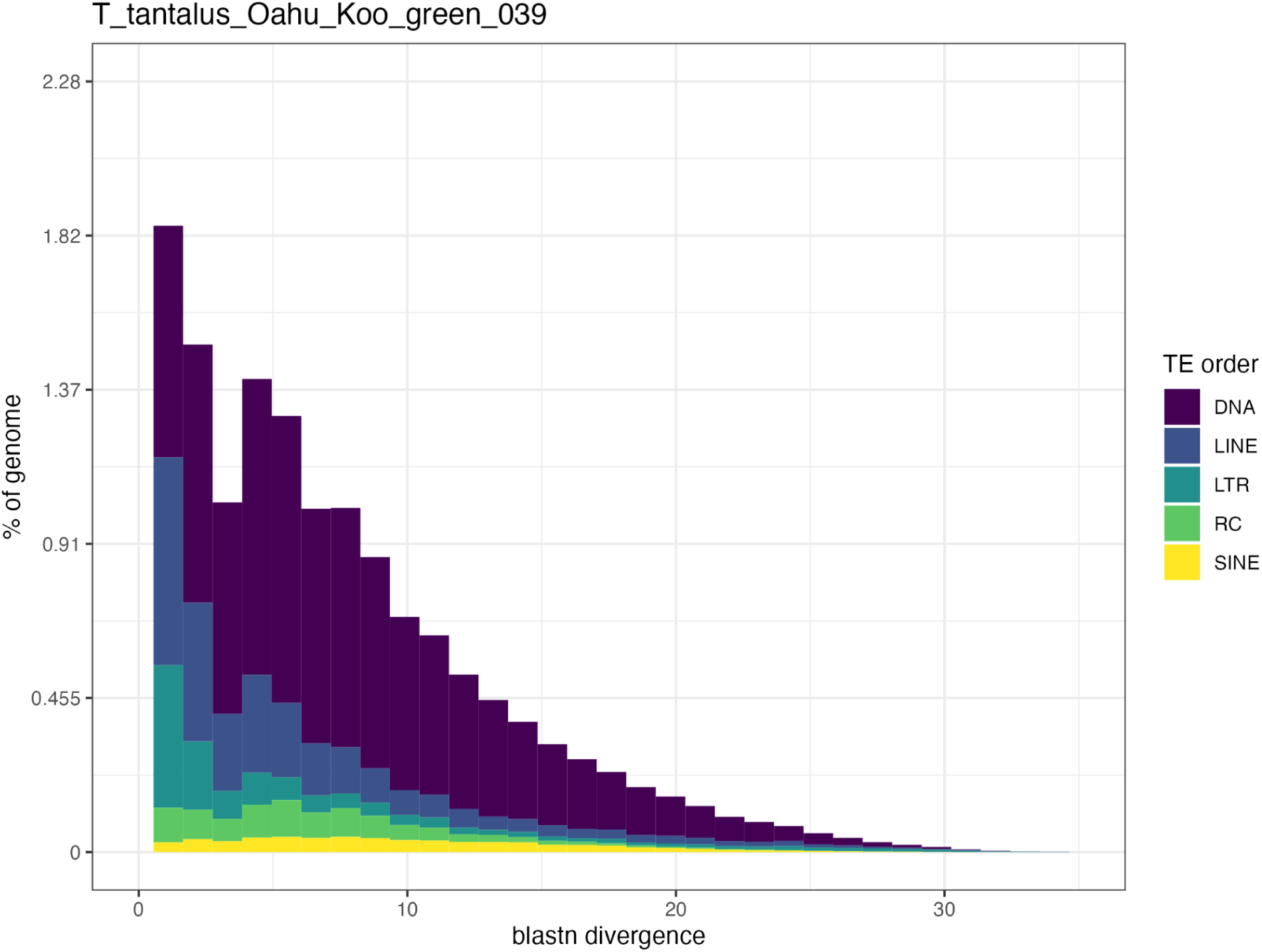
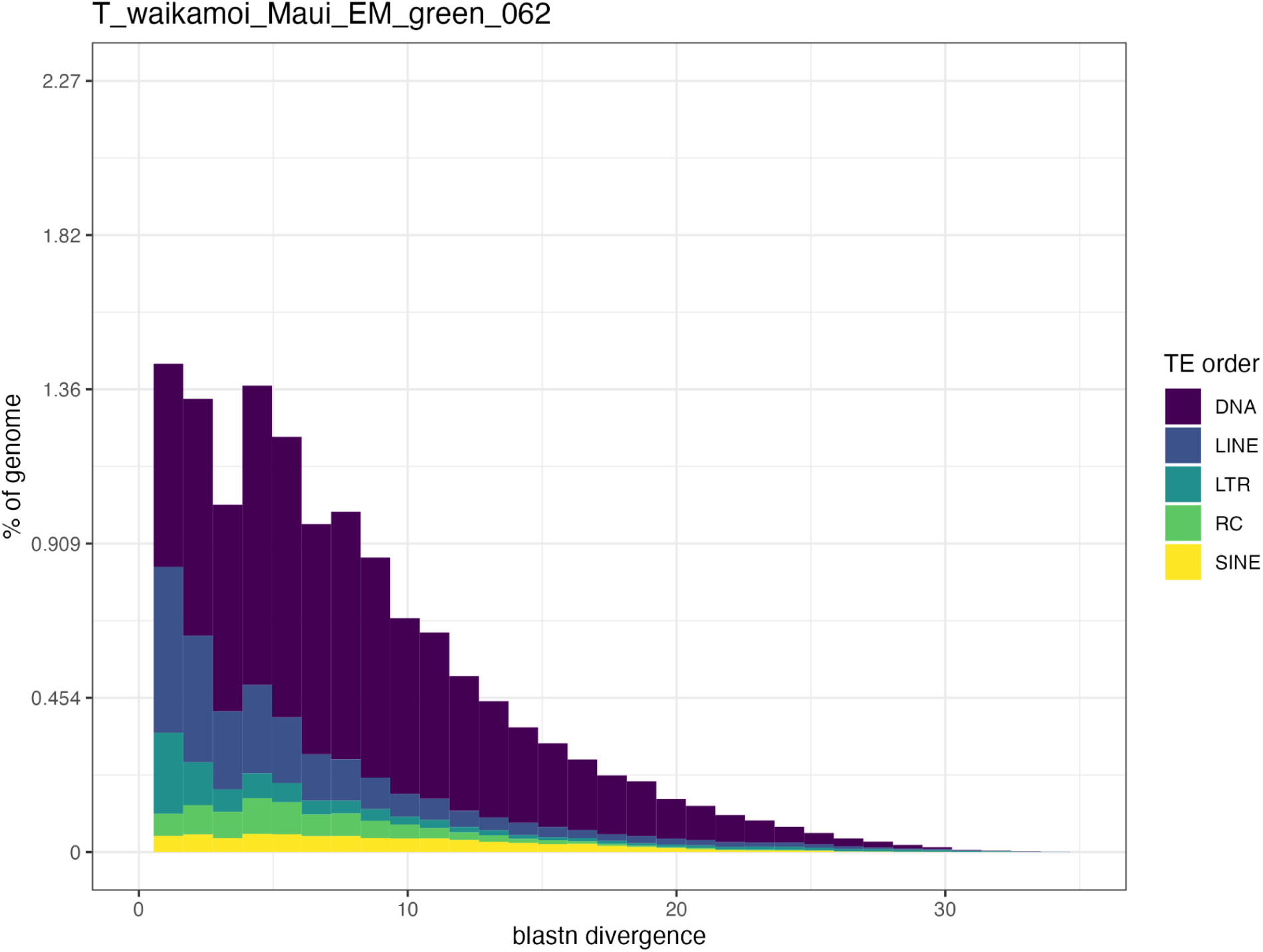
Repeat landscape plots for all *Tetragnatha* individuals. The x-axis represents the percent divergence of each TE’s consensus sequence using blastn divergence. The y-axis represents the percent of the genome per bin of percent divergence. Each color is an order of TEs (DNA, LINE, LTR, RC, SINE).

**Supplementary figure 04.**
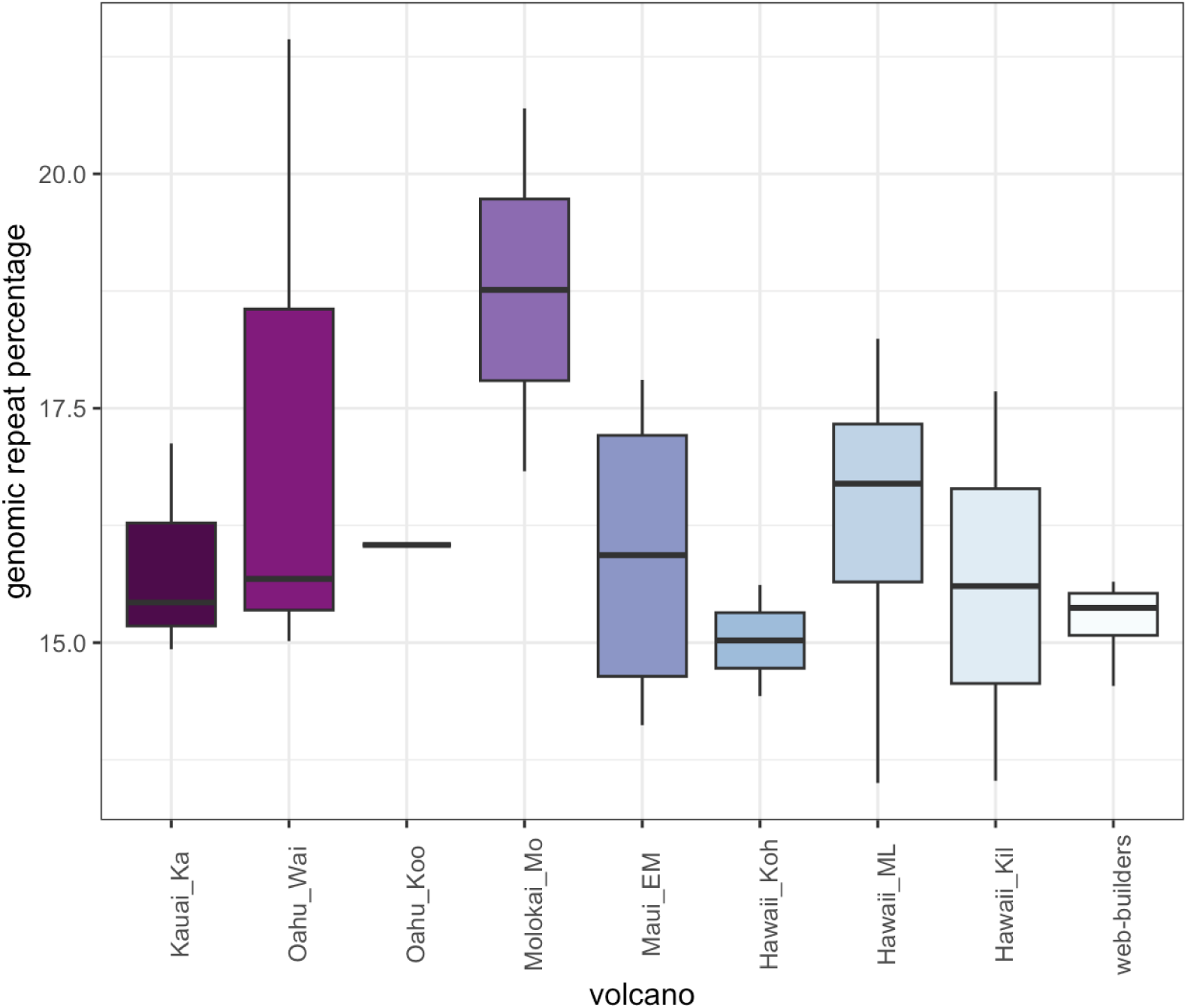
Genomic repeat content by volcano. Spiny-leg species on multiple volcanoes were included on their respective volcanoes.

**Supplementary figure 05.**
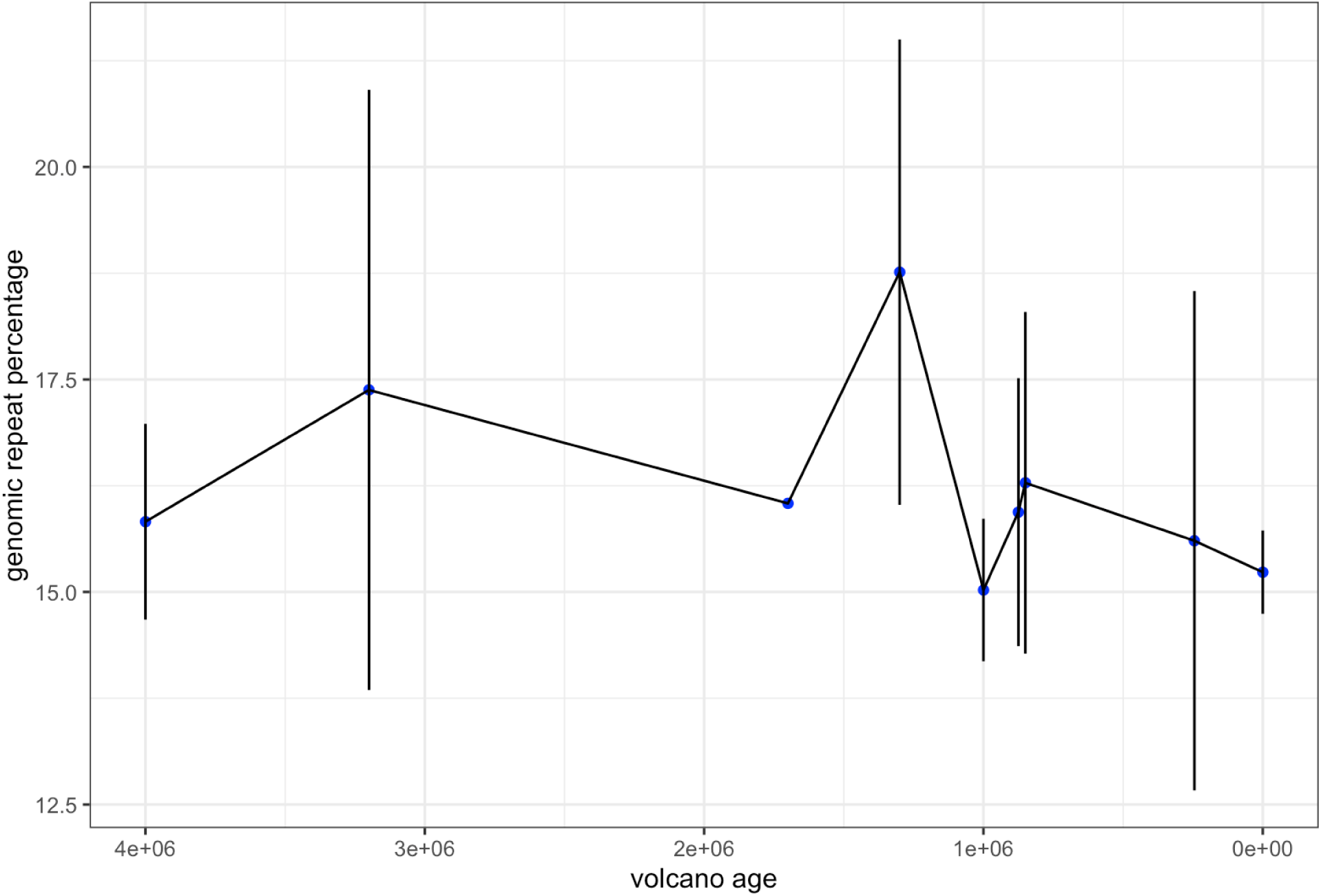
**Relationship between average repeat content and volcanic age.**

**Supplementary figure 06.**
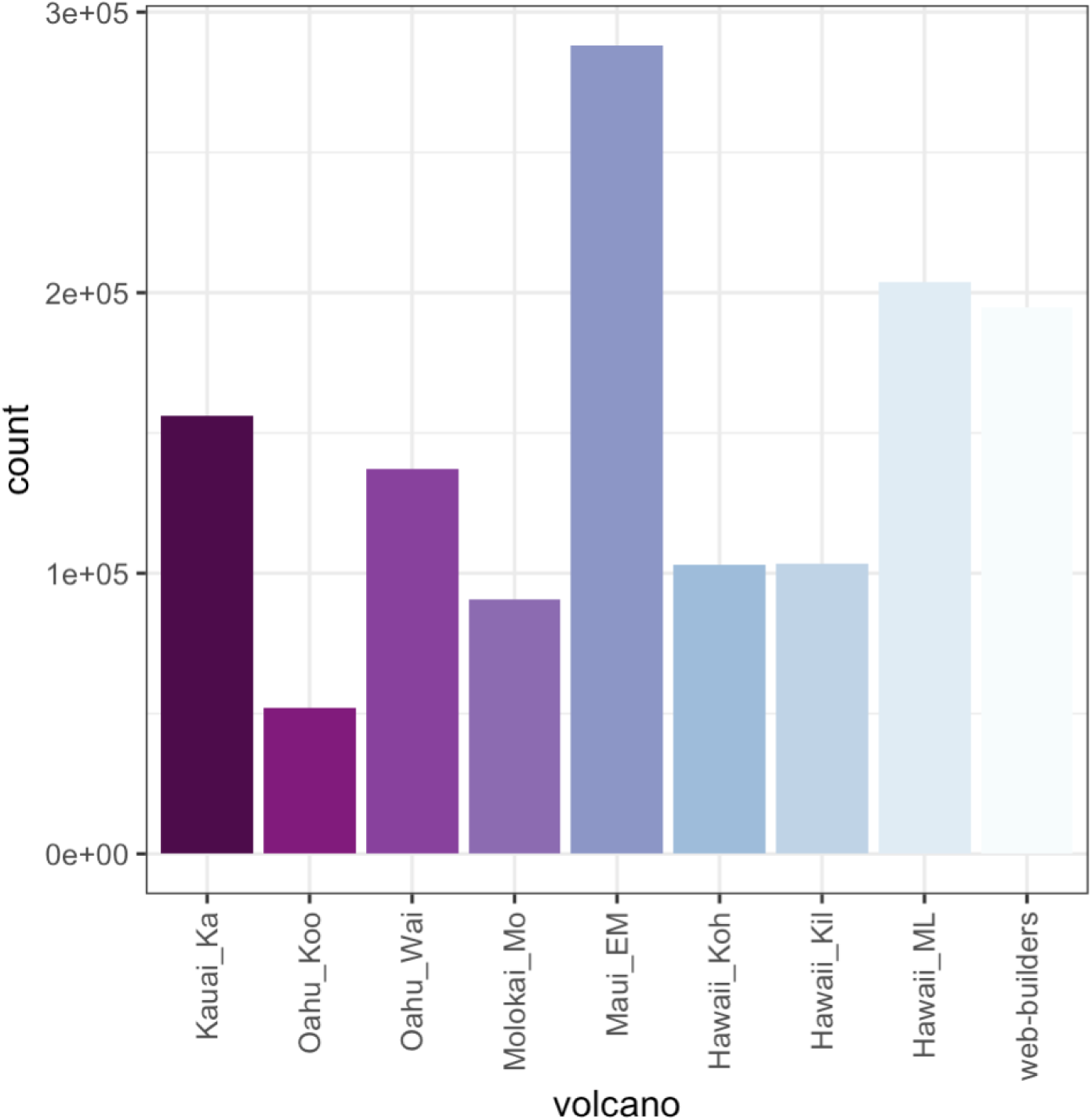
**Average abundance of DNA/hAT transposons across volcanoes.**

